# Transcriptional evaluation of the developmental accuracy, reproducibility and robustness of kidney organoids derived from human pluripotent stem cells

**DOI:** 10.1101/238428

**Authors:** Belinda Phipson, Pei X Er, Lorna Hale, David Yen, Kynan Lawlor, Minoru Takasato, Jane Sun, Ernst Wolvetang, Alicia Oshlack, Melissa H Little

**Author notes:** Corresponding author: Professor Melissa H. Little, NHMRC Senior Principal Research Fellow, Murdoch Children’s Research Institute, Flemington Rd, Parkville, 3052, Melbourne, VIC, Australia.

## Abstract

We have previously reported a protocol for the directed differentiation of human induced pluripotent stem cells to kidney organoids comprised of nephrons, proximal and distal epithelium, vasculature and surrounding interstitial elements. The utility of this protocol for applications such as disease modelling will rely implicitly on the developmental accuracy of the model, technical robustness of the protocol and transferability between iPSC lines. Here we report extensive transcriptional analyses of the sources of variation across the timecourse of differentiation from pluripotency to complete kidney organoid, focussing on repeated differentiations to day 18 organoid. Individual organoids generated within the same differentiation experiment show Spearman’s correlation coefficients of >0.99. The greatest source of variation was seen between experimental batch, with the enrichment for genes that also varied temporally between day 10 and day 25 organoids implicating nephron maturation as contributing to transcriptional variance between individual differentiation experiments. A morphological analysis revealed a transition from renal vesicle to capillary loop stage nephrons across the same time period. Distinct iPSC clones were also shown to display congruent transcriptional programs with inter-experimental and inter-clonal variation most strongly associated with nephron patterning. Even epithelial cells isolated from organoids showed transcriptional alignment with total organoids of the same day of differentiation. This data provides a framework for managing experimental variation, thereby increasing the utility of this approach for personalised medicine and functional genomics.

## Introduction

The analysis of genetic disease at a molecular level involves comparisons between primary cell types derived from affected and unaffected individuals. Such approaches face many challenges, not least of all the capacity for the cell type being examined to reflect the disease phenotype. The ability to derive induced pluripotent stem cells (iPSC) from any somatic cell of a patient (Takahashi et al. 2007), together with directed differentiation protocols to generate specific target cell populations, has provided a capacity to specifically model the cell types affected by the disease. This has proven effective in the case of cardiomyocytes and neurons for the validation of causative genes and the analysis of the specific effect on cellular function (Bellin et al. 2013; Kim et al. 2013; Phelan et al. 2016; Ardhanareeswaran et al. 2017; Aksoy et al. 2017). The utility of such approaches has substantially improved in the case of monogenic conditions with the advent of CRISPR editing by facilitating the creation of isogenic corrected lines for direct comparison with mutant lines (Jang & Ye 2016; Paquet et al. 2016; Howden et al. 2017).

Recent iPSC differentiation protocols have been developed in which multiple component cell types arise and self-organise in a fashion similar to the developing tissue (Ader & Tanaka 2014; Huch & Koo 2015). Protocols have now been reported for the generation of optic cup, pituitary cerebral cortex, intestine and stomach organoids (Suga et al. 2011; Eiraku et al. 2011; Spence et al. 2011; Nakano et al. 2012; Lancaster et al. 2013; Kadoshima et al. 2013; McCracken et al. 2014). Unlike protocols generating a relatively homogeneous target cell type, such as cardiomyocyte or neuron, the cellular complexity of such structures increases the likelihood of substantial variation between individual experiments. This will be compounded when comparing distinct cell lines for disease modelling, even if these represent patient and isogenic control lines, as individual iPSC clones show variation in rates of cell proliferation and response to ligand that may not be associated with an inherited genetic defect. Indeed, organoid protocols also tend to involve many weeks of differentiation to reach such complex endpoints, with any variation in the differentiation process across that time period also contributing to experimental variation. Understanding the robustness and reproducibility of such differentiation protocols is therefore paramount.

We have recently developed a stepwise protocol for the generation of kidney organoids from human pluripotent stem cells (Takasato et al. 2015; Takasato et al. 2016). During this differentiation protocol, we have attempted to move the cells in a stepwise manner through the developmental milestones required for kidney morphogenesis during normal fetal development. Using a monolayer culture format, an initial induction of the primitive streak is achieved via canonical Wnt signalling, intermediate mesoderm patterning via FGF9 signalling, and then a 3D aggregate is formed with nephrogenesis induced via a pulse of canonical Wnt signalling (Takasato et al. 2016). The resulting organoids contain >9 distinct cell types, representing the key epithelial cell types of the forming filtration units, the nephrons (podocytes, parietal epithelium, proximal tubule, loop of Henle, distal tubule), together with collecting ducts, endothelium, pericytes and interstitial fibroblasts. While this differentiation protocol was based upon our understanding of mouse development, the presence of key morphological structures positive for common markers in mouse development supports the assumption of strong developmental congruence between mouse and human kidney development. However, with such a large number of distinct cell types present, the possibility for simple variation in the relative proportions of each component cell type raises the challenge of experimental reproducibility or the capacity for identifying novel disease-associated transcriptional change.

In this study, we provide a more complete transcriptional and morphological evaluation of the protocol for generation of kidney organoids from human pluripotent cell lines. Using RNA sequencing (RNA-seq) to globally profile gene expression of whole kidney organoids, we examine the degree of variation between organoids within a single differentiation experiment, between different differentiation experiments, between different iPSC clones and from the epithelial compartment isolated from within organoids. Focussing on a single timepoint of differentiation (day 18), we show striking transcriptional correlation between all organoids analysed (r^2^>0.956), with the highest congruence between individual organoids within the same differentiation experiment (r^2^>0.997) and the lowest congruence (r^2^>0.923) resulting from batch effects likely to reflect media components, RNA extraction and library preparation. A rank list of the most variable genes aligns closely with temporal gene expression changes across the organoid differentiation protocol, suggesting that these transcriptional variations reflect differences in relative organoid maturation, particularly with respect to nephron morphogenesis. Profiling of LTL-enriched or EpCAM-enriched epithelial populations from within organoids showed strong alignment with total organoids of the same day of differentiation, suggesting that this approach to reducing cellular complexity continues to reflect stage of development. Finally, a comparison of organoids generated using genetically distinct control iPSC cell lines showed no greater variation between lines than between batches. In total, these analyses provide a bioinformatic framework for approaching disease modelling using such a complex model, highlighting the need to minimise batch to batch variation when generating organoids from distinct lines.

## Results

### Temporal characterisation of *in vitro* human kidney organoid differentiation demonstrates an appropriate embryogenic differentiation program

The protocol previously developed for the production of kidney organoids commences with a single vial of single cell-adapted human iPSC plated onto matrigel, with this representing day -1. At Day 0, differentiation is commenced within fully defined APEL media together with a specific regime of small molecule and recombinant proteins across a total period of 25 days, at which point approximately 20-30 individual organoids are generated (Takasato et al. 2016). We refer to this as one differentiation experiment. An initial intra-experimental evaluation, performed as a comparison between individual organoids from the point of aggregation (day 7) using RNAseq analysis, has previously been reported (Takasato et al. 2015). This included the profiling of three replicate individual kidney organoids collected at the time of aggregation at day 7, +3 days, +11 days and +18 days (referred to here as day 7, 10, 18 and 25) (GEO accession GSE70101). Here, we retrospectively extended this dataset by collecting triplicate RNA samples at the beginning of culture (day 0, pluripotent stem cells at plating) and day 4 (end of CHIR induction). In this case, the triplicates examined represented individual wells from the same starting vial. The examined timepoints within this complete differentiation protocol are referred to as days 0, 4, 7, 10, 18 and 25 (Figure 1A). See ‘Methods’ for information regarding sample and library preparation for RNA-Sequencing.

**Figure 1.**
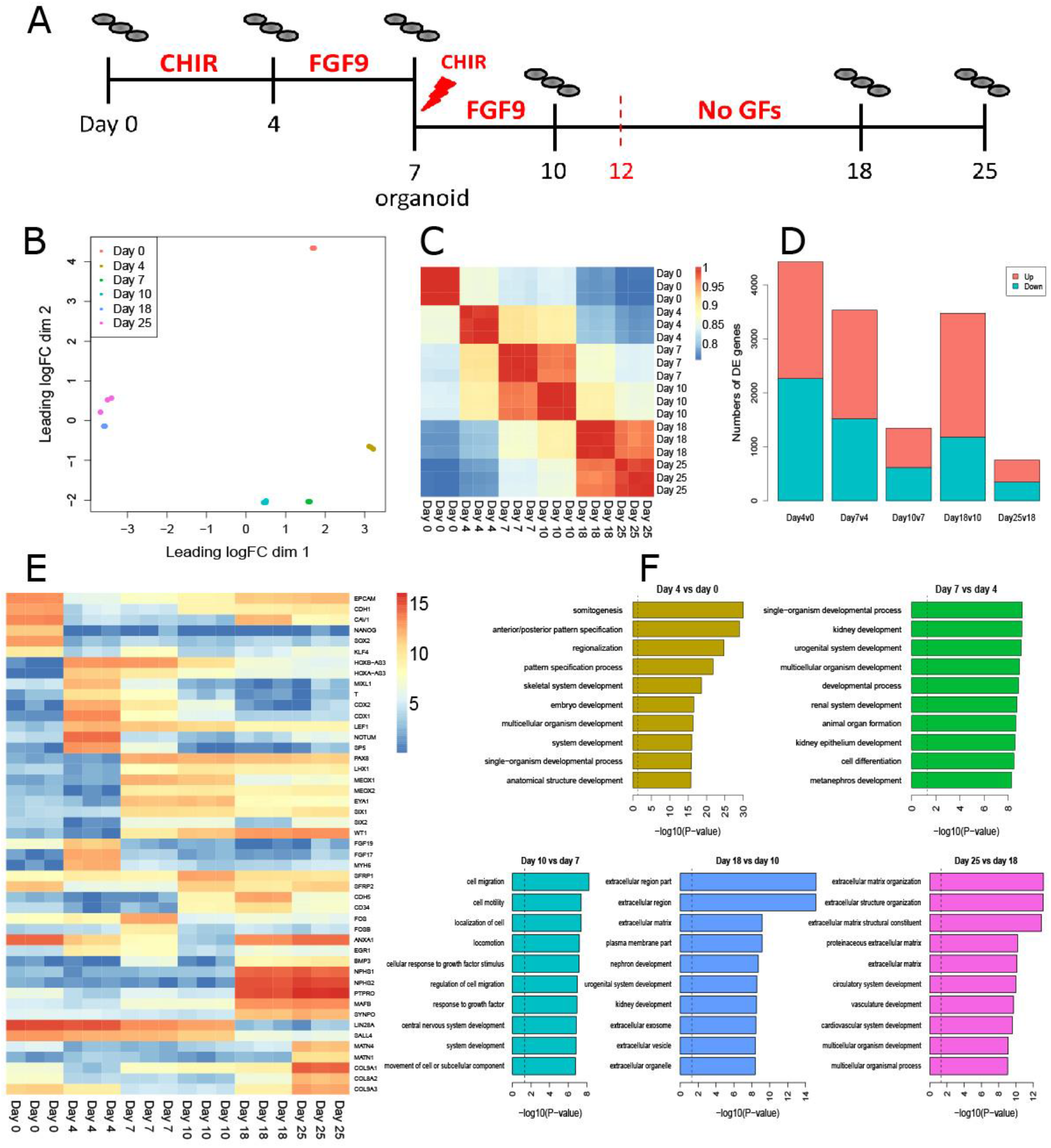
Temporal characterisation of *in vitro* human kidney organoid differentiation demonstrates an appropriate embryogenic differentiation program. **A.** Diagram of the differentiation protocol to show the timepoints collected. Material collected from day 0 and day 4 were individual wells from a 6 well plate of monolayer cultures. **B.** Multi-dimensional scaling plot of all samples collected across the timecourse demonstrates a clear developmental trajectory. **C.** Correlation plot between samples across the timecourse showing tight correlation within individual replicates together with strong similarity between day 7 and 10 and day 18 and 25. **D.** Number of significant differentially expressed genes up and downregulated between consecutive timepoints of differentiation. **E.** Heatmap of the most highly differentially expressed genes between each timepoint. **F.** GO analysis of differentially expressed genes between consecutive timepoints.

A multidimensional scaling plot based on the 500 most variable genes revealed very high congruence (Spearman correlation > 0.986) between all samples at each timepoint showing high reproducibility between organoids within a given differentiation experiment (Figure 1B,C). Two approaches were used to investigate the molecular changes occurring across the differentiation protocol; i) consecutive pair-wise gene expression comparisons to find those genes increasing or decreasing most between each timepoint and ii) unsupervised clustering of genes across time points to identify genes displaying synexpression.

While substantial changes in gene expression occurred between each consecutive timepoint (Figure 1D), these were most pronounced at the beginning of the protocol (induction of primitive streak), between day 4 and day 7 (formation of intermediate mesoderm), and day 10 and day 18 (kidney differentiation). The transition from day 0 to 4 (2160 genes up, 2267 genes down, |logFC| > 1, FDR < 0.05; Supplementary Table 1) mimics the induction of primitive streak, with early loss of epithelial markers (e.g. EPCAM, CDH1 and CAV1) and pluripotency markers (e.g. NANOG, SOX2, KLF4 and OCT4), with a concurrent increase in expression of posterior primitive streak markers (e.g. T, MIXL1, CDX1 and CDX2) and HOX genes (Figure 1E). WNT signalling pathways were strongly up-regulated at day 4 (p=1.96e-07, Supplementary Table 2) due to the addition of a GSK3beta inhibitor, CHIR99201, with strong up-regulation of the transcription factor SP5 (logFC = 12, FDR =1.16e-306), which is directly regulated via canonical WNT signalling (Fujimura et al. 2007). Enriched GO terms for the top 100 up-regulated genes between day 4 and day 0 included somitogenesis, anterior/posterior pattern specification and regionalisation (Figure 1F, Supplementary Table 2).

From day 4 to 7 (2008 genes up, 1528 genes down, |logFC| > 1, FDR < 0.05; Supplementary Table 3), with the removal of CHIR and the addition of FGF9, the patterning was assumed to represent intermediate mesoderm. As anticipated between these timepoints, down-regulated genes were enriched in WNT signalling (p=1.76e-08), with NOTUM (modulates WNT signalling pathways) and SP5 significantly down-regulated. PAX8 and LHX1, which are essential for intermediate mesoderm induction (Tsang et al. 2000; Bouchard et al. 2002; Wang et al. 2005), were significantly up-regulated across this time period while markers present in metanephric mesenchyme and ultimately in the nephrogenic mesenchyme, such as MEOX1/2, EYA1, SIX1/2, WT1 (Brunskill et al. 2008), were also strongly up-regulated by day 7 (all logFC > 6.8). The most down-regulated genes between day 4 and 7 included primitive streak / definitive mesoderm markers CDX1, MIXL1 and T as well as FGF19, FGF17 and MYH6. GO terms involved in urogenital system development, kidney development, renal system development, and kidney epithelium development are significantly over-represented for the top 100 up-regulated genes between day 4 and day 7 (Figure 1F, Supplementary Table 2).

Between day 7 and 10, there were fewer genes differentially expressed (731 up-regulated, 617 significantly down-regulated, |logFC| > 1, FDR < 0.05, Supplementary Table 4). Up-regulated genes include secreted frizzled-related proteins (SFRP1/2), cadherins (CDH5, CDH1) and CD34; while down-regulated genes included FOS, FOSB, ANXA1 (annexin A1), HOXA-AS3, HOXB-AS3, KLF4, EGR1, BMP3. There was also a further down-regulation of CDX1/2, MIXL1, T and SP5. GO terms significantly over-represented for up-regulated genes included cell migration, cell motility and cellular response to growth factor stimulus (Figure 1F, Supplementary Table 2). While the upregulation of CDH1 might suggest the commencement of epithelial structure formation, overall genes associated with epithelium development were over-represented in the down-regulated genes indicating that at this time point there is little evidence of nephron induction. Between day 10 and day 18 there was a large up-regulation of genes suggestive of vascular differentiation and nephron development (2292 up-regulated genes, 1185 down-regulated genes, |logFC| > 1, FDR < 0.05, Supplementary Table 5). Indeed, at day 18, one of the most highly expressed genes in the organoids included markers of the visceral epithelial cell type, the podocyte, a key cellular component of the forming glomerulus. Such genes included NPHS1, NPHS2, PTPRO, MAFB, SYNPO, ITIH5, CLIC5, NPNT, all of which are expressed in podocytes (Brunskill et al. 2011). Simultaneous with the upregulation of nephron markers between day 10 and 18 was the downregulation of markers of progenitor mesenchyme, including LIN28A, MEOX1, CITED1 and EYA1. GO terms over-represented for the top 100 up-regulated genes include nephron development, urogenital system development and kidney development (Figure 1F, Supplementary Table 2).

Fewer transcriptional changes were seen between day 18 and day 25 (403 up-regulated, 359 significantly down-regulated, |logFC| > 1, FDR < 0.05, Supplementary Table 6), however GO terms related to extracellular matrix were strongly over-represented for the top 100 up-regulated genes (Figure 1F, Supplementary Table 2). For the top 100 down-regulated genes, top enriched GO terms included vasculature development, cardiovascular system development, circulatory system development and blood vessel development, although some of the genes in these GO terms were also up-regulated (Supplementary Table 2).

Our second approach to the analysis of the molecular program across the entire time series was to look for synexpression clusters. This analysis allowed for clusters of genes to be identified with non-linear expression patterns across the time course. Genes were selected that showed expression changes across any time point across the time course (ranked by F-statistic, |logFC|>1). Fuzzy clustering was then performed allowing for 20 expression clusters to be identified (Supplementary Figure 1), and core contributing genes with high membership scores (>0.5) for each cluster were extracted (Supplementary Table 3). Clusters of interest are show in Figure 2A. PCR for selected genes within clusters 5, 6, 7, 10, 14, 16 and 17 validated these patterns of gene expression in independent differentiations (Figure 2B).

**Figure 2.**
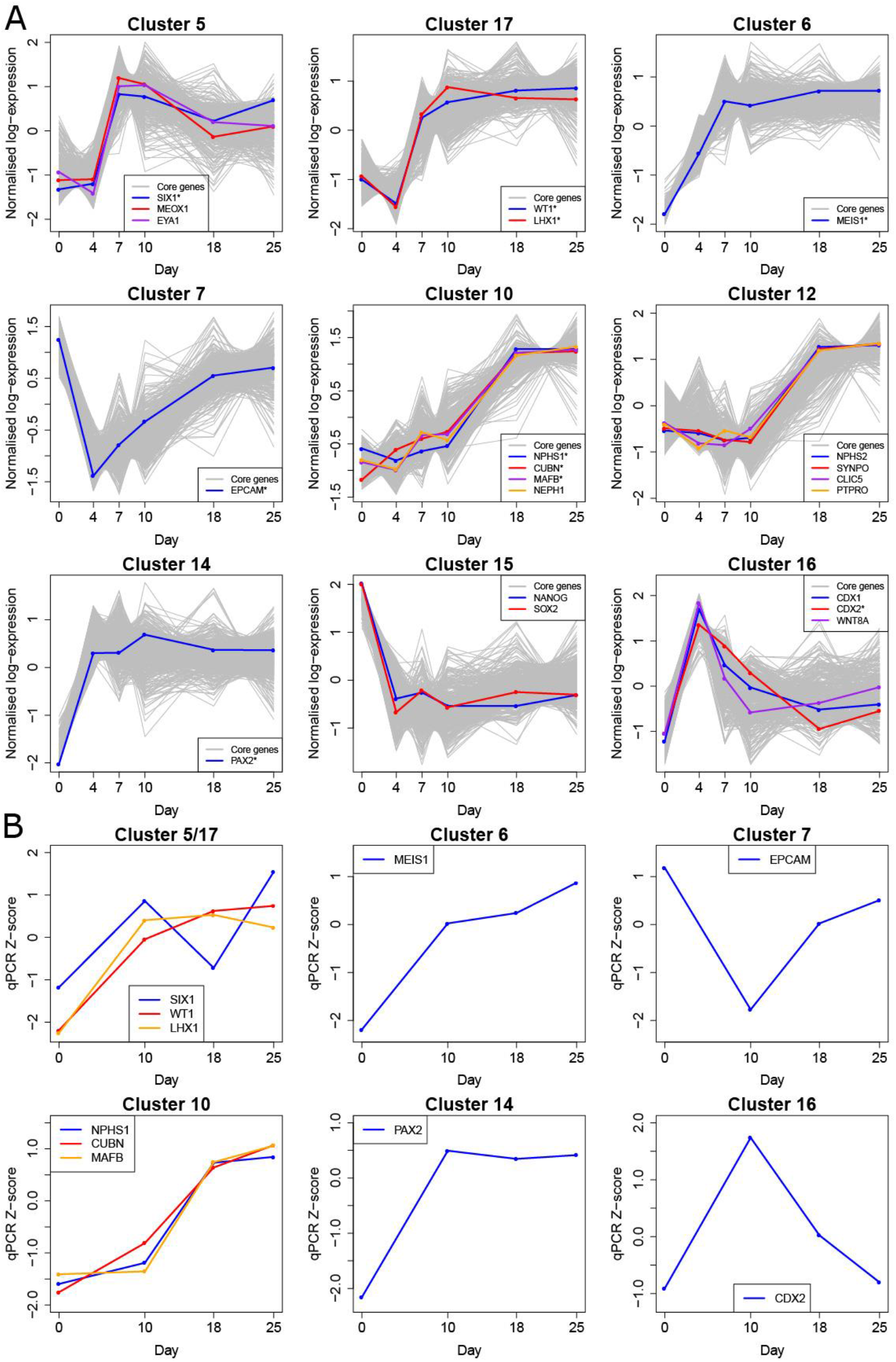
Analysis of the molecular program of human kidney differentiation. **A.** Selected cluster plots showing synexpression across the developmental time. Core genes with membership scores > 0.5 are shown in grey. Example genes of interest are shown in colour for each cluster. Genes with an asterisk were validated with qPCR. Lists of core genes for each cluster are seen in Supplementary Table 3. **B.** qPCR validation of selected genes from clusters 5, 17, 6, 7, 10, 14 and 16 in independent samples across days 0, 10, 18 and 25 of the differentiation protocol.

In many instances, core genes in a given cluster are easily associated with anticipated profiles across a presumptive stepwise differentiation from pluripotency to kidney. For example, cluster 15 genes (including NANOG and SOX2) were highly expressed at day 0 and subsequently lost expression from day 4 onwards, indicative of a loss of pluripotency (Figure 2A). Cluster 7 included loss of expression of epithelial genes, such as EPCAM, as the pluripotent stem cells began differentiation (days 0 to 4) which then returned as nephrons arose within the organoids (Figure 2A). Genes in cluster 14 were activated from day 4 onwards, and included PAX2, an early marker of intermediate mesoderm (Figure 2A). Cluster 6 contains genes up-regulated from day 7 onwards, and included MEIS1, a marker of renal stroma. Cluster 16 genes, including CDX1 and WNT8A, were upregulated exclusively at day 4 with core genes enriched for Wnt signalling pathway genes (p-value=0.00012, modified hypergeometric test). Clusters 5 and 17 displayed upregulation from day 7 onwards and included LHX1 (intermediate mesoderm marker), as well as markers of the cap mesenchyme (MEOX1, EYA1, SIX1, WT1). Finally, clusters 10 and 12 contain genes that are turned on in maturing nephrons from day 18, such as the podocyte (MAFB, SYNPO, PTPRO, NPHS2, KIRREL/NEPH1 and CLIC5) and proximal tubule (CUBN) markers. Selected genes from these clusters were validated by qPCR in independent samples at days 0, 10, 18 and 25 of the differentiation protocol (Figure 2B). The top 20 enriched GO and KEGG terms for the core genes in each cluster are listed in Supplementary Table 4. Key transcription factors, identified as such through the FANTOM5 consortium (http://fantom.gsc.riken.jp/5/sstar/Browse_Transcription_Factors_hg19), were identified within the core genes for each cluster, however only a subset of these transcription factors had a defined JASPAR motif (Table 1, Supplementary Table 5). These included MYCN in cluster 3, MAFB and HNF1A in cluster 10, NR3C1 and IRF2 in cluster 11, PAX2 in cluster 14, SOX2, FOXD3, INSM1 and SPIB in cluster 15, EBF1, NR2F1 and HNF1B in cluster 17 and TCF3 in cluster 19. This dataset will facilitate analysis of transcriptional networks driving synexpression within these clusters.

**Table 1:** Core genes that are also transcription factors in the 20 synexpression clusters.

| Cluster | Num core genes | Num FANTOM5 TFs | TFs with JASPAR motif | Num hits |
| --- | --- | --- | --- | --- |
| 1 | 220 | 12 |  |  |
| 2 | 70 | 1 |  |  |
| 3 | 77 | 11 | MYCN |  |
| 4 | 79 | 16 |  |  |
| 5 | 107 | 23 |  |  |
| 6 | 206 | 39 |  |  |
| 7 | 142 | 8 |  |  |
| 8 | 88 | 3 |  |  |
| 9 | 123 | 5 |  |  |
| 10 | 217 | 10 | MAFB, HNF1A |  |
| 11 | 131 | 8 | NR3C1, IRF2 |  |
| 12 | 162 | 6 |  |  |
| 13 | 168 | 11 |  |  |
| 14 | 207 | 31 | PAX2 |  |
| 15 | 156 | 17 | SOX2, FOXD3, INSM1, SPIB |  |
| 16 | 106 | 7 |  |  |
| 17 | 158 | 32 | EBF1, NR2F1, HNF1B |  |
| 18 | 167 | 7 |  |  |
| 19 | 299 | 23 | TCF3 |  |
| 20 | 92 | 11 |  |  |

### Analysis of the source of transcriptional variation between organoids within and between experiments

The cellular complexity present within an individual kidney organoid is likely to result in significant technical and biological variation. This is a particular challenge for disease modelling where variation unassociated with the disease state may exist. Sources of variation can arise from biological variation intrinsic to cell line or clone and technical variation resulting from vial / passage, batch variation in culture components and variations in downstream processing procedures. While correlation between organoids generated within the initial differentiation experiment was very high (r^2^>0.986), we wanted to understand the level of transcriptional variability that could arise when differentiation experiments are repeated. Using the same cell line as used in the time series RNAseq study, CRL1502-C32, organoids (18 in total) were generated from another 6 differentiation experiments with RNA collected from individual organoids at day 18 of culture (Figure 3A). These distinct differentiation experiments were also performed across a total of 12 months and hence across different reagent batches (culture media, recombinant growth factors). Organoids from experiments 3, 4 and 5, while all commencing with a distinct vial of starting iPSC, were all differentiated in parallel, and hence were classed as one “batch” (Figure 3A). RNA isolation, library preparation and sequencing were also performed at the same time for these three experiments. The remaining organoids from experiments 1, 2, 6 and 7 were differentiated at different times, and considered to be distinct batches. All samples were sequenced at the same facility using the same protocol (see Methods).

**Figure 3.**
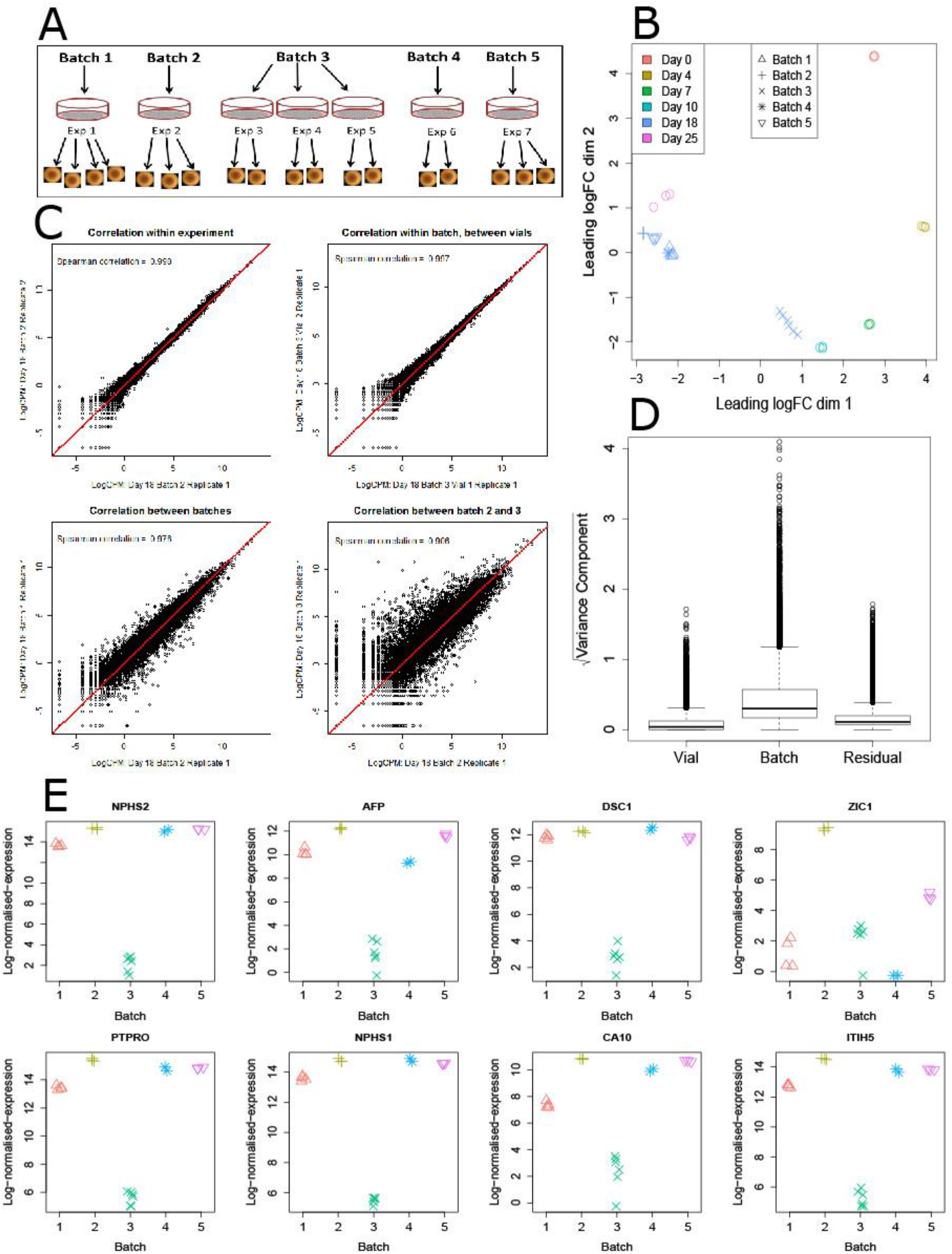
Analysis of the source of transcriptional variation between organoids within and between experiments. **A.** Diagram of all day 18 organoids profiled showing the relationship between batch, experiment and organoid. Each batch represented a single starting vial of undifferentiated iPSC cells. Each experiment represented a single differentiation from starting cells using the same protocol. Batches were separated in time but experiments within a single batch were performed at the same time. Within batch 3, three differentiation experiments were performed concurrently from three separate vials. Samples were individual organoids from within an experiment. **B.** Multi-dimensional scaling plot of all day 18 organoids as well as the time series samples indicating batch, day and individual sample. All day 18 organoids were positioned between day 10 and day 25 organoids. **C.** Scatterplots of log-normalised-expression for pairs of representative samples showing the degree of correlation between organoids. Organoids differentiated within an experiment were incredibly highly correlated, as were organoids differentiated between vials in batch 3. Correlations between organoids from different batches were slightly lower, ignoring organoids from batch 3. Organoids from batch 3 had the lowest correlation with organoids from other batches. All pairwise correlations shown in Supplementary Figure 2. **D.** For each gene, an analysis partitioning the source of variation was performed. Boxplots show the contribution to each source of variation (batch, vial and residual) across all genes. The largest contribution to variability is the batch, with vial-to-vial and residual variation contributing much less to the total variation observed. **E.** Top 8 most variable genes, stratified by batch. Except for ZIC1, the most variable genes show a strikingly similar pattern of expression, with organoids in batch 3 have lower expression of these genes compared to the remaining batches. Heatmap of expression of top 50 most variable genes in Supplementary Figure 3 and full table of variability analysis results in Supplementary Table 6.

Day 18 organoids differentiated in the same batch were highly concordant, with tight clustering shown on a multi-dimensional scaling plot that included the time course data (Figure 3B). Overall, the transcriptional correlation between all day 18 organoids generated across all batches was also very high (average Spearman’s correlation = 0.956), with the highest correlation observed between organoids differentiated at the same time (average Spearman’s correlation = 0.997, Figure 3C, Supplementary Figure 2). Organoids differentiated in batch 3 tended to be slightly less correlated with organoids generated from the remaining batches (average Spearman’s correlation = 0.923, Figure 3C, Supplementary Figure 2). When compared with the time course experiment, all day 18 organoids were positioned between day 10 and day 25 on a multi-dimensional scaling plot, with batch 3 organoids closer to day 10 organoids compared to the remaining organoids (Figure 3B).

Our experimental design allowed us to explore three levels of variability; (i) batch to batch variability, which captures variation arising due to performing differentiation experiments at different points in time; (ii) vial to vial variability for organoids differentiated in parallel in batch 3; and finally (iii) organoid to organoid variability and all other unknown sources of variation, referred to here as residual variation. For each gene, a random effects analysis was performed which allowed us to obtain an estimate of the contribution of these three sources of variability, referred to here as variance components (Supplementary Table 6). Across all genes, the largest contribution to the transcriptional variability observed was batch to batch variability, with vial to vial variability only a small contributor and residual varance being minimal (Figure 3D).

Genes with the largest total variability between batch were examined in order to gain an understanding of how the variability in the transcriptional profiles between the day 18 organoids may arise. Many of the top most variable genes were related to nephron maturation (NPHS2, PTPRO, NPHS1) with these genes following a strikingly similar expression pattern across the batches (Figure 3E). Examining the top 50 most variable genes from the day 18 organoids (Supplementary Figure 3) across the original time course data showed a highly consistent temporal expression pattern, with low/no expression from day 0 to day 10, followed by a dramatic increase in expression coincident with nephron formation (from day 10 onwards) (Figure 4A). In line with this, the top 50 most variable genes were enriched in synexpression clusters 10 and 12 (7 genes in cluster 10, 16 in cluster 12, Supplementary Figure 4).

**Figure 4.**
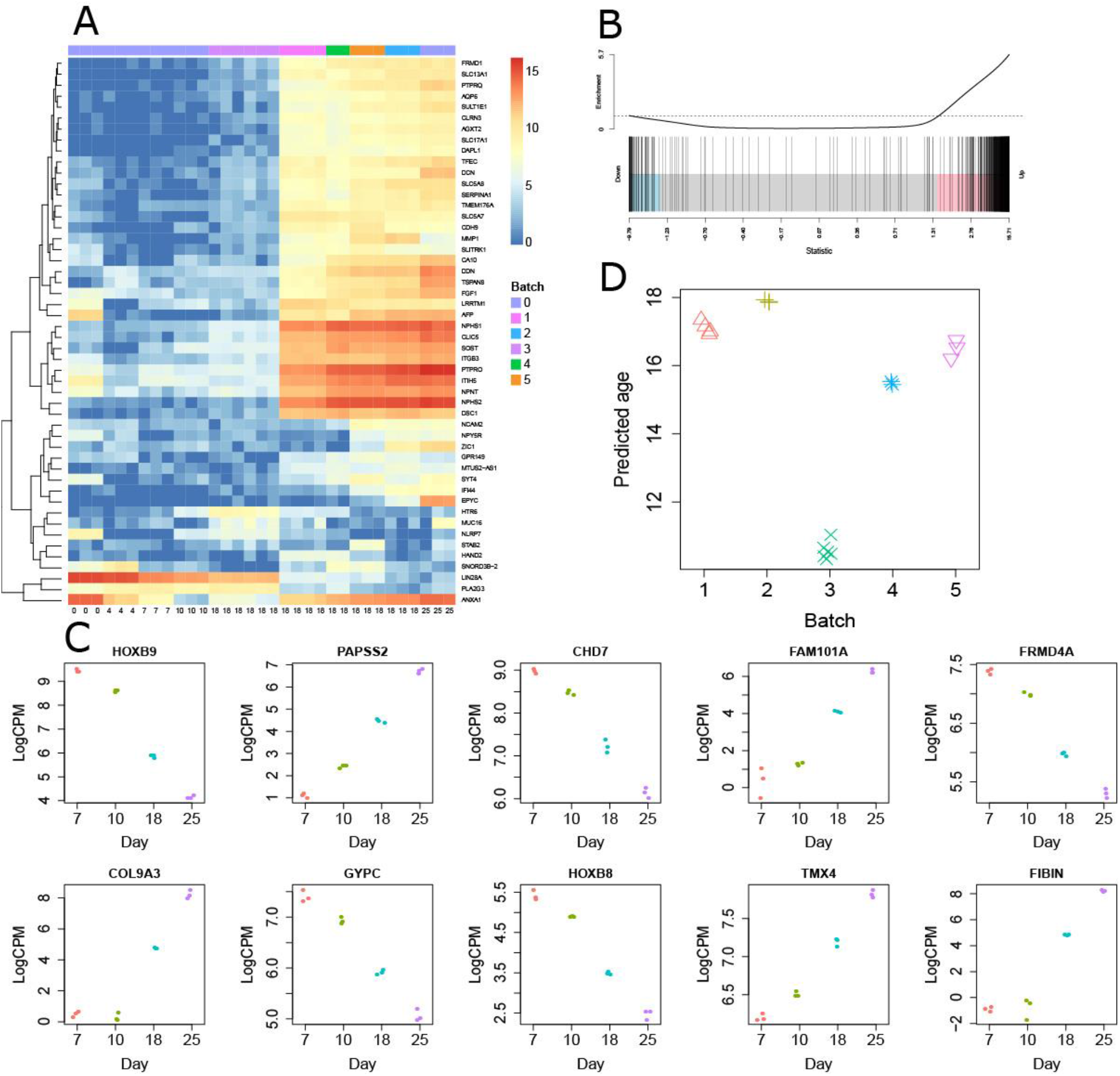
Analysis of the association between differentiation and transcriptional variability so as to predict a relative degree of organoid maturation between batches. **A.** Analysis of the temporal pattern of expression of genes most highly variable between batches of kidney organoids at day 18.The majority of the genes show a highly consistent pattern of expression across the time series. **B.** ROAST analysis suggesting association between differentiation and transcriptional variability. Highly variable genes at day 18 are strongly enriched between days 25 and 10 of the time course. **C.** Identification of key genes that predict relative maturation. **D.** Prediction of organoid age based upon key genes marking differentiation progression.

In order to test whether the highly variable genes were related to nephron maturation, we performed differential expression testing between days 10 and 25 in the original time course data to indicate nephron formation related changes. The top 500 most highly variable genes between organoids at day 18 were strongly enriched for these nephron-related genes, with approximately 80% of the variable genes significantly up-regulated between day 10 and day 25 (p-value = 0.0005, ROAST test, Figure 4B). 10 genes were identified that were most linearly associated with a developmental progression from day 7 to day 25 of differentiation (Figure 4C). Using these 10 genes, a multivariate linear regression was constructed for the day 7 to 25 time series data to obtain regression coefficients. Defining the additional day 18 organoids as new sample data, the normalised “age” of any given organoid relative to the time series data was calculated (Figure 4D). This analysis mirrored the trend seen in the multi-dimensional scaling plots, with batch 3 organoids having the youngest predicted ages (10.3-11 days) and the remaining organoids batches (1, 2, 4 and 5) closer in developmental age to day 18 (median predicted ages: batch 1 = 17.1, batch 2 = 17.9, batch 4 = 15.5, batch 5 = 16.5). This approach may be applicable for normalisation of variation between kidney organoid batches for a given iPSC line.

### Morphological and transcriptional evidence for nephron patterning in organoids between day 10 and day 25

The morphology of the forming nephron has been well documented in mouse (Little et al. 2007; Harding et al. 2011) with the definition of distinct stages of nephron patterning and segmentation defined (Figure 5A). Nephrons arise via a mesenchymal to epithelial transition as an initial pretubular aggregate (PTA) forms an epithelial renal vesicle (RV; Stage I nephron) which almost immediately displays definitive distal (*JAG1*, *LHX1*) and proximal (*WT1*) gene expression (Georgas et al. 2009) (Figure 5A). These structures undergo elongation to form comma-shaped (CSB) and then S-shaped bodies (SSB; Stage II nephrons). The SSB show expression of distinct medial segment markers with *JAG1* expression becoming medial (Georgas et al. 2009; Lindström et al. 2015). Involution of the proximal end of the elongating nephron to form the glomerular cup into which endothelial and mesangial cells migrate represents the capillary-loop stage (CL; Stage III) nephron (Georgas et al. 2009) (Figure 5A). In the Trimester 1 human kidney, very similar structures have been described (O’Brien et al. 2016). However, in both mouse and human, these arise at stereotypical locations adjacent to a dichotomously branching ureteric epithelium. Using co immunofluorescence for distal (CDH1), medial (JAG1) and proximal (WT1/NPHS1) protein markers, we examined the structures present within human kidney organoids from day 10 to 25 for evidence of normal patterning and segmentation (Figure 5B). No evidence for epithelial structures was present before day 10 (Figure 5D), at which time PTA and RV became readily observable (Figure 5BD). This was followed by a subsequent elongation and segmentation of epithelial units until, by day 18, most structures represented late-SSB to CL (Figure 5BC). JAG1 protein clearly moved from the distal RV to the medial SSB / CL nephron, as has been described in mouse (Lindström et al. 2015). This analysis suggests that the formation of a nephron from PTA to CL takes approximately 6 to 8 days, as has been described in the developing human kidney (O’Brien et al. 2016). *In vivo*, the collecting duct epithelium is present before nephrons arise and the forming nephrons invade the collecting duct tip to form a contiguous epithelium (Georgas et al, 2009). Notably, a detailed evaluation of the GATA3+ECAD+ epithelium suggests that this arises in association with the forming nephrons rather than representing a distinct epithelial structure which fuses with the nephrons (Figure 5D). Indeed, some forming epithelial structures appear to have branching structures which are GATA3+ECAD+ epithelium that may represent the distal invading tip unit (Figure 5E). Unlike the developing organ *in vivo*, there was little evidence of continued nephron formation in these kidney organoids. Rather, the initial induction events appeared to give rise to early nephrons that then mature.

**Figure 5.**
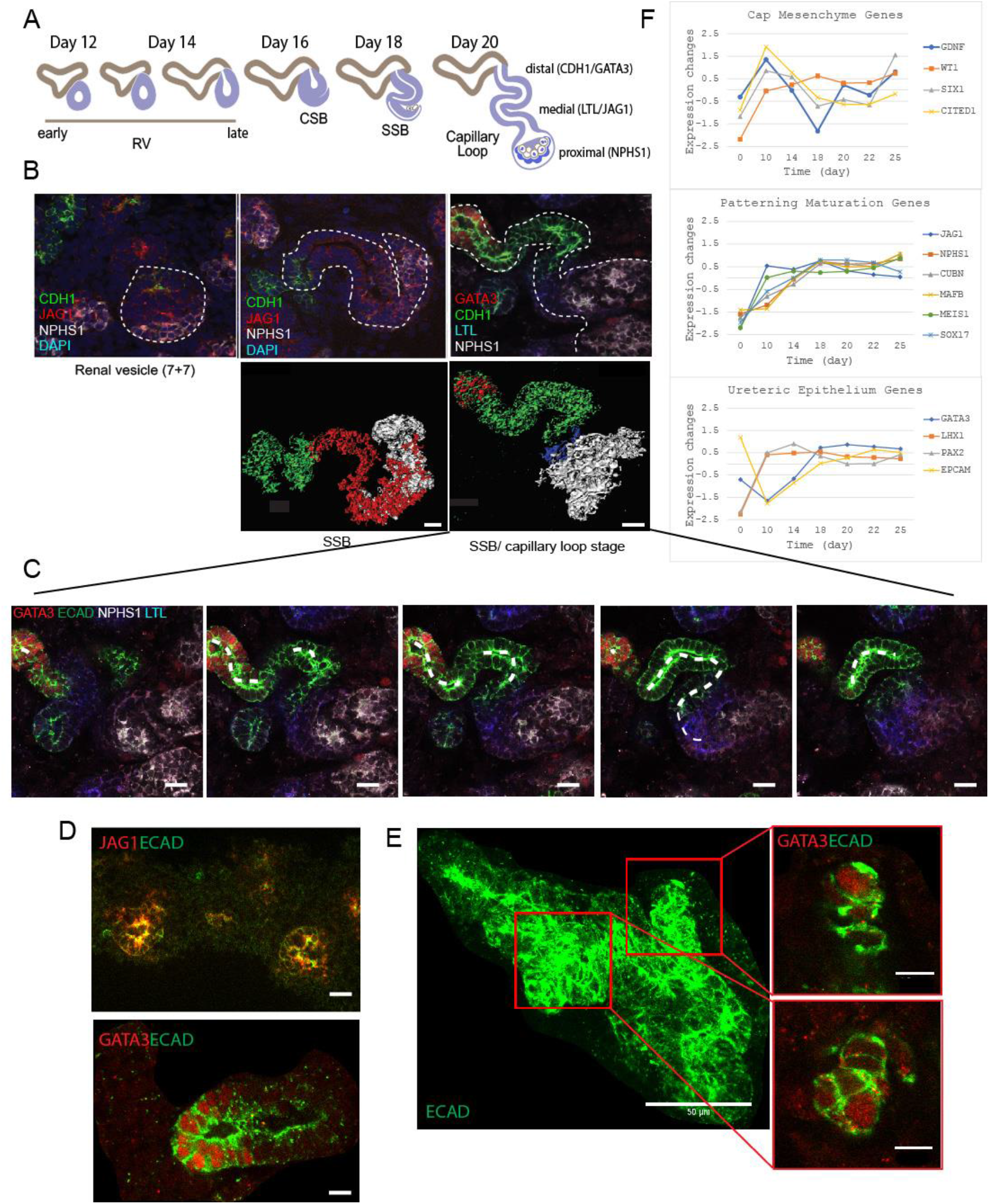
Evidence for nephron patterning and segmentation to capillary loop stage. **A.** Diagram of anticipated morphological changes across nephron formation in mouse kidney. **B.** Immunofluorescence of nephrons forming within organoids across time using markers for proximal nephron (white; NPHS1), distal RV / medial nephron (red; JAG1) and distal nephron (green; CDH1). **C.** Serial single Z slices through confocal images of a single capillary-loop stage nephron showing segmentation along the length of the tubule and evidence for a lumen passing in and out of the plane of the image. Immunofluorescence performed using antibodies to markers of collecting duct (red; GATA3), epithelium (green; CDH1) and proximal nephron (white; NPHS1). Scale bar = 10uM. **D.** High resolution images of early nephrons arising at Day 10 of culture showing staining for JAG1 (red) and ECAD (green) (scale bar = 10uM) or GATA3 (red) and ECAD (green) (scale bar -15uM). **E.** Maximal intensity projection (MIP) of an ECAD+ (green) epithelial structure arising at Day 14 with single Z slices through two side branches revealing evidence for GATA3+ECAD+‘tip’ structures. Scale bar = 50uM for MIP and 10uM for insets. **F.** QPCR of key nephrogenic genes between day 10 and 25 shows gradual loss of nephrogenic progenitors from day 10 to 14 and formation of nephrons commencing at the same time.

Overall, this morphological analysis supports the concept that substantial transcriptional variation between day 10 and day 25 is in large part due to variation in relative nephron maturation or nephron number. In order to analyse this in a more qualitative fashion, RNA was collected from day 14, 18, 20, 22 for qPCR of core genes implicated in nephron maturation (cluster 10 and 12), mesenchyme-to-epithelium transition (MET) (cluster 7), nephrogenesis (cluster 11), renal interstitium development (cluster 13), nephron commitment (cluster 17) and renal fate specification (cluster 20) (Figure 2A). This analysis showed an increase in nephron and stromal gene expression with time and a concomitant reduction in metanephric mesenchymal markers (Figure 5F), again supporting the hypothesis of gradual nephron patterning and maturation across this period of time.

### Correlation between kidney organoids generated using distinct iPSC clones

All the data presented thus far was collected from differentiations performed using a single human iPSC clone, CRL1502-C32 (Briggs et al. 2013). This line was derived from fetal fibroblasts collected from a normal female using footprint-free episomal reprogramming (Yu et al, 2009). To examine the experimental variation arising from a distinct iPSC clone with a different genome, kidney organoids generated from another female control human iPSC line, RG_0019.0149.C6, were also generated. The resulting organoids displayed a similar morphology to CRL1502-C32 at the level of immunofluorescence (Supplementary Figure 5). RNA-seq profiling of organoids from this line was performed on two day 18 organoids from each of three separate but simultaneous differentiation experiments (6 organoids in total). In addition, triplicate day 0 and day 7 cultures of RG_0019.0149.C6 were profiled, one from each differentiation experiment. Combining organoids from both RG_0019.0149.C6 and CRL1502-C32 revealed striking concordance across the differentiation protocol, with organoids clustering based on differentiation duration and not cell line (Figure 6A). Day 18 organoids from RG_0019.0149.C6 clustered between CRL1502-C32 batch 3 and the remaining batches (Figure 6A). Spearman’s correlation for each pairwise organoid across the time series and across the two cell lines again showed that organoids at the same time point were more highly correlated than organoids within each cell line between the time points (Figure 6B). This confirmed that batch effects are higher contributors to variation than genetic background or individual iPSC clone. All differentiations using this second line represented a single batch as all were differentiated simultaneously. It was possible, however, to examine gene-wise variation arising from distinct vials of the same line as well as any residual (kidney-kidney plus unknown) variation at day 18. Organoids grown from the same vials of RG_0019.0149.C6 were highly correlated (Figure 6C, mean Spearman’s correlation = 0.998), with this correlation remaining high between organoids grown from different vials of the same clone (Figure 6C, mean Spearman’s correlation = 0.997). As for the CRL1502-C32 organoids, a random effects analysis showed that the vial contribution was relatively small across all genes, with the residual variation contributing more transcriptional variability in general (Figure 6E, Supplementary Table 7). However, the most highly variable genes could be attributed to vial-to-vial variability, with differences driven by organoids between vial 2 and vial 3 (Supplementary Figure 6). Variable genes with high vial-vial contributions were over-represented in immune-related pathways, while variable genes with high residual contributions were over-represented in signalling receptor activity (Supplementary Figure 7, Supplementary Table 8).

**Figure 6.**
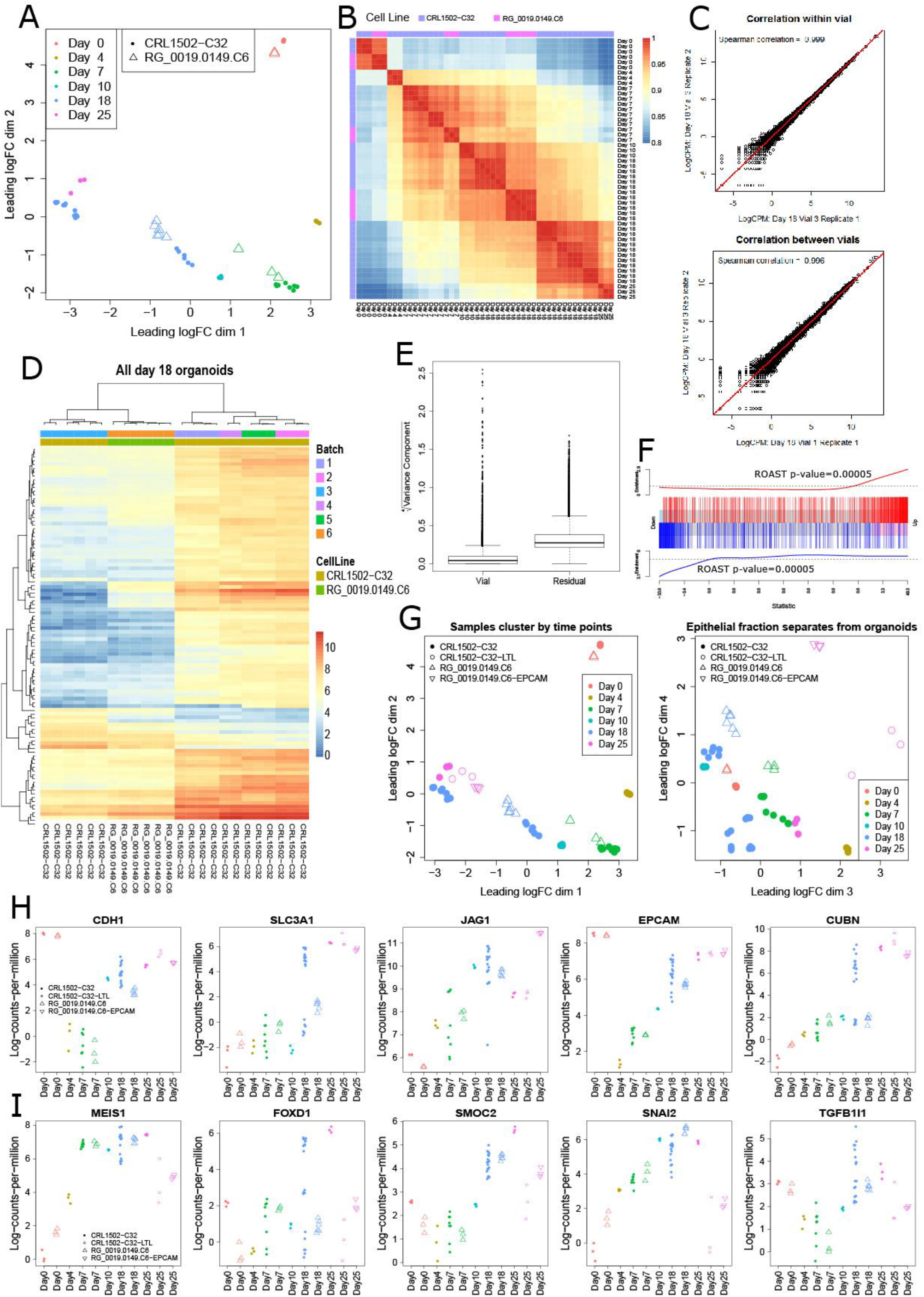
Transcriptional variation between iPSC lines and temporal concordance between total organoid and enriched nephron epithelium. **A.** Multi-dimensional scaling plot for total organoids generated from two distinct cell lines, CRL1502-C32 and RG_0019.0149.C6, across the entire differentiation protocol. Day 0 (3 replicates), 7 (3 replicates) and 18 (6 replicates across 3 vials of starting material). **B.** Heatmap of the Spearman’s correlation between all RG_0019.0149.C6 and CRL1502-C32 organoids. Organoids correlate more closely based upon stage of differentiation than cell line across the time points. **C.** Scatterplots of log-normalised-expression for pairs of RG_0019.0149.C6 organoids illustrates mean Spearman’s correlation between organoids generated from the same (0.998) or different starting vials (0.997). All day 18 RG_0019.0149.C6 organoids were grown concurrently and hence represent one batch. **D.** Heatmap with unsupervised hierarchical clustering of log-counts-per-million showing the top 100 differentially expressed genes between day 18 and day 10 in the original time course data. All day 18 organoids from the two cell lines are included. Batch 6 contains organoids from the RG_0019.0149.C6 cell line. All other batches as previously defined. A clear trend in expression of these maturity-related genes indicates that the RG_0019.0149.C6 organoids are more mature than CRL1502-C32 batch 3 organoids, but have not reached the same developmental maturity as organoids in batches 1, 2, 4 and 5. **E.** Random effects modelling estimated two variance components for each gene; vial-to-vial variability and residual variability. Boxplots show contributions from the two sources of variation genome-wide. Similarly to the analysis of CRL1502-C32 organoids, vial-vial variability was a minor contributor to total variability genome-wide. **F.** Barcodeplot showing enrichment of genes differentially expressed between CRL1502-C32 day 18 batch 3 organoids and RG_0019.0149.C6 day 18 organoids when compared to day 18 versus day 10 CRL1502-C32 organoids. This indicates that genes differentially expressed between the cell lines are enriched for maturity-related genes, rather than genes specific to the genotype of the cell lines. **G.** Multidimensional scaling plots of day 25 enriched nephron epithelium samples from the two cell lines, labelled CRL1502-C32-LTL and RG_0019.0149.C6-EPCAM, compared to all other total organoid samples. The total organoids and enriched samples cluster by time point in dimension 1 and 2, however in the lower dimensions 3 and 4 there is a separation between the total organoids and enriched nephron epithelium samples. **H.** Log-normalised-expression of 5 epithelial-related genes for all total organoids and enriched nephron epithelium samples showing high expression of these genes in the nephron epithelium samples. **I.** Log-normalised-expression of 5 interstitial-related genes for all total organoids and enriched nephron epithelium samples showing reduction in expression of these genes in the nephron epithelium samples.

The differences between the day 18 organoids from the two different cell lines could again be associated with the relative degree of maturation of the organoids, rather than genotype differences. We defined a set of maturity-related genes by performing differential expression analysis between day 18 and day 10 of the original time course data. Performing unsupervised clustering of all day 18 organoids using the top 100 maturity related genes indicated that RG_0019.0149.C6 day 18 organoids were “older” than batch 3 CRL1502-C32 day 18 organoids, but “younger” than the remaining CRL1502-C32 organoids (batches 1, 2, 4 and 5; Figure 6D). In addition, we performed a differential expression analysis between RG_0019.0149.C6 day 18 organoids and CRL1502-C32 day 18 organoids from Batch 3 that were generated with the same replication design. This gave us a set of up- and down-regulated genes differentially expressed between the two cell lines at day 18. These genes were highly enriched in the set of maturity-related genes (Figure 6F, ROAST p-values = 0.0005).

### Epithelial-enriched populations isolated from organoids cluster with age-matched total organoids

The variation in global transcriptional profile between individual experiments even within the same cell line is likely to be exacerbated by the diversity of cell types present. In order to simplify the pool of cells being analysed, we also performed MACs-enrichment of the EpCAM+ epithelial fraction of organoids generated from the RG_0019.0149.C6 cell line, and the LTL+ epithelial fraction from the CRL1502-C32 cell line, both at day 25 of organoid differentiation. Multidimensional scaling analysis across the first two dimensions showed that both of the epithelial-enriched profiles clustered alongside the global organoid profile of the same differentiation time point, illustrating the contribution of this cell type to the global profile (Figure 6G). Plotting the lower dimensions showed separation of total organoid and epithelium-enriched samples (Figure 6G). Plotting the log-normalised expression of a set of selected epithelial and stromal markers confirmed high expression of epithelial-enriched profiles (Figure 6H) and depleted expression of interstitial markers in both epithelium-enriched populations (Figure 6I). The LTL fraction showed distinctly higher expression of JAG1 consistent with the selective enrichment of the proximal tubular epithelium (LTL+) versus all epithelium (EpCAM+). While this approach will not overcome inherent differences in nephron maturation between batches, it does reduce the complexity of the transcriptional profile potentially improving the capacity to identify cell-type specific transcriptional changes. Such an approach is likely to assist in disease modelling.

## Discussion

The prospect of applying organoids to disease modelling or the analysis of morphogenesis *in vitro* rests implicitly on the accuracy of the model and the robustness and reproducibility of the protocol within and between lines. This becomes an even greater challenge with the cellular complexity of an organoid versus a relatively homogeneous cellular endpoint. Kidney organoids represent arguably the most complex human organoid generated to date. With this increased cellular complexity comes challenges in terms of variation in degree of cell patterning or relative proportions of cell types between experiments.

Despite this challenge, in this study we report transcriptional profiling of 51 kidney organoids, including whole and fractionated samples, and show a remarkable degree of congruence in profiles between organoids generated across considerable technical and biological variation. Overall, this analysis revealed that transcriptional variation is greatest between experimental batch. This strongly implicates technical variables that may include changes in batch of media, growth factors, RNA isolation reagent and cDNA library preparation. An analysis of the most variable genes between batch showed strong congruence with temporally variable genes associated with the period of nephron formation. We hypothesise that this results from variation in relative maturation between differentiation experiments, even using the same cell line. However, it is also possible that this reflects differences in the segmentation of individual nephrons within a given organoid and/or variations in the relative amounts of nephron versus non-nephron cell types in different organoids. Indeed, there appears to be greater variation between distinct organoid batches than between different starting cell lines, implying that the greatest source of variation is in the reagents used during culture. Hence, variation at day 18 may also reflect subtle shifts in patterning across the entire differentiation.

While the robustness of the kidney organoid protocol would suggest that disease modelling may be feasible, it also identifies the challenges of comparisons performed between individual patient and control cell lines. Analysis of the sources of variation has allowed us to identify a set of 10 genes that facilitate a retrospective estimation of relative organoid ‘age’ for any given experiment. As this is a *post hoc* analysis, this does not ensure that lines to be compared will all reach the same level of maturation. It is possible to exclude the most variable genes from any transcriptional comparison, however this may also remove information of relevance to the phenotype of a particular line. Another approach to reducing technical variation is to select for specific cell types from within the organoid. As a proof of concept, we report here the profiling of epithelial cells enriched from organoid cultures via magnetic sorting for two different markers of the epithelial phenotype. While we could show that these were enriched for epithelial markers, and depleted for stroma, they still aligned most closely with total organoids matched for differentiation stage with the organoids from which the epithelium was sourced. Therefore, analysis of selected cellular components from within organoids is likely to improve comparative phenotyping.

As the greatest source of variation appears to arise from differences in technical parameters, rather than cell line, cellular compartment or individual organoid, this would suggest that any comparisons between patient and control lines must be performed concurrently with as many technical variables controlled as possible. To assist in normalisation, we provide a list of genes that are most variable between batches at a single timepoint. Utilising this list of variable genes can also be performed to generate an estimate of relative organoid maturity, but this does not provide a solution if disease versus control lines repeatedly show differences in relative maturation state. Despite these challenges, we have successfully identified disease-related transcriptional changes between a patient and CRISPR-edited isogenic control line by removing from the analysis the 500 most variable genes identified in this study (Forbes et al, submitted).

In summary, this study demonstrates that directed differentiation of human iPSC to kidney organoid is robust, reproducible and transferable between different pluripotent stem cell lines. This is the first detailed analysis of its type of the sources of transcriptional variation in a complex multicellular organoid model. The implications of this study are significant with respect to what this variation means for the design of disease modelling studies. As the greatest source of variation appears to arise from differences in technical parameters, rather than cell line, cellular compartment or individual organoid, this would suggest that any comparisons between patient and control lines must be performed concurrently with as many technical variables controlled as possible.

## Methods

### Human pluripotent stem cell derivation and directed differentiation to kidney organoids

Human induced pluripotent stem cells were generated using Sendai reprogramming as previously described. CRL1502.C32 is a female human iPSC line derived from ATCC CRL-1502 fetal fibroblasts (Briggs et al. 2013). RG_0019.0149.C6 is a female human iPSC line derived from fresh skin fibroblasts harvested from an adult. The protocol for directed differentiation to kidney organoid has been previously described (Takasato et al. 2015; Takasato et al. 2016).

### RNA sequencing, data acquisition, alignment and QC

RNA from 51 kidney organoid differentiations was extracted using the QIAGEN RNeasy micro kit and sequencing libraries prepared using the standard Illumina protocols. For epithelial fractions, these were obtained from pooled organoids from the same differentiation. All samples were sequenced at the Institute for Molecular Bioscience in Queensland, Australia. The STAR aligner (v2.4.0h1) (Dobin et al. 2013) was used to map the 75bp single end reads to the human reference genome (hg19) in the two pass mapping mode. Uniquely mapped reads were summarised across genes with feature Counts (v1.4.6) (Liao et al. 2014) using Gencode Release 19 comprehensive annotation. Subsequent analyses of the count data were performed in the R statistical programming language using the Bioconductor (Gentleman et al. 2004) packages edgeR (Robinson et al. 2010), limma (Ritchie et al. 2015), RUVSeq (Risso et al. 2014), Mfuzz (Futschik & Carlisle 2005), the annotation package org.Hs.eg.db; as well as the lme4 R package (Bates et al. 2015). Highly expressed genes were defined as having at least one count per million in at least two or three samples and were retained for statistical analysis. The threshold for selecting highly expressed genes is determined by the minimum group sample size in the dataset being analysed. In addition, genes encoding for ribosomal protein, mitochondrial genes, pseudogenes genes and genes without annotation were removed prior to TMM normalisation (Robinson & Oshlack 2010) and statistical analysis.

### Differential expression analysis of time series data

The time series RNA-Seq data consists of three replicates at each of six time points: Day 0, Day 4, Day 7, Day 10, Day 18 and Day 25. The samples for Days 7, 10, 18 and 25 were obtained and sequenced together, with additional data for the earlier time points, Days 0 and 4, generated at a later date. In order to account for possible batch effects, the differential expression analysis was performed using RUVSeq in conjunction with edgeR. RUVSeq was performed using empirical control genes, identified as the 5000 genes that varied the least across the time course data, based on an F-statistic. Genes that were differentially expressed at consecutive time points were identified as those that had an adjusted false discovery rate of less than 5%. Genes significantly differentially expressed with an absolute logFC of at least one and false discovery rate less than 5% were identified with a TREAT analysis (McCarthy & Smyth 2009). Gene ontology testing was performed using the goana function in limma, adjusting for gene length bias (Young et al. 2010), as well as the web-based ToppGene suite (Chen et al. 2009).

### Multidimensional scaling plots and unsupervised clustering

For all datasets, multidimensional scaling plots based on the top 500 most variable genes were used to visualise the greatest sources of variation, specifying pairwise distance metrics in the limma package. Genes that displayed similar patterns of expression across the time course data were clustered using fuzzy c-means clustering. The clustering was limited to genes that showed evidence of differential expression across the time course based on an F-statistic, and with an absolute logFC of at least one between at least one comparison. For this analysis, each time point was compared to the remaining time points. This identified 7682 genes to use as input for the Mfuzz algorithm. The counts were transformed to log counts per million, adding a small offset of 0.25 proportional to the library sizes. Each of the three replicates were averaged per time point, and the data standardised such that each gene had a mean of zero and standard deviation of one to ensure genes with similar changes in expression are close in Euclidean space. The soft clustering approach assigned each gene gradual degrees of membership, ranging from zero to one, to each of the 20 clusters. A core set of genes for each cluster was identified by specifying a cut-off on the membership score of 0.5, with numbers of core genes per cluster ranging from 70 to 299. Gene ontology and KEGG pathway analysis on the core genes was performed using the goana and kegga function in limma, adjusting for gene length bias (Young et al. 2010), and further explored using the ToppGene suite.

### Transcription factor analysis

Transcription factors for hg19 were downloaded from the FANTOM5 website (http://fantom.gsc.riken.jp/5/sstar/Browse_Transcription_Factors_hg19). The R packages JASPAR2016 and TFBSTools (Tan & Lenhard 2016) were used to explore binding site hits for selected motifs in the core genes for each of the fuzzy clusters.

### Random effects modelling

In order to study the different variance components when comparing across experiments and batches a multi-level random effects model was fitted to the Day 18 organoid data for each cell line separately using the lmer function in the lme4 R package. The data was transformed to log counts per million adding a small prior count of 0.5 before model fitting, and variance components for experiment, vial and residual extracted for the CRL1502-C32 organoids, and vial and residual components extracted for the RG_0019.0149.C6 organoids. Genes were ranked according to total variation, which was obtained by summing the variance components. For each gene, the greatest contributor to the variability was obtained by calculating proportions of each variance component to the total variation.

### Bright Field Imaging and Immunofluorescence Imaging of cultures

Bright field images were taken using the Nikon TS-1000 inverted microscope. For the kidney organoid, antibody staining was performed as described previously (Takasato et al., 2016). The following antibodies and dilutions were used: mouse anti-ECAD (1:300, 610181, BD Biosciences), goat anti-GATA3 (1:300, AF2605, R&D Systems), sheep anti-NPHS1 (1:300, AF4269, R&D Systems), LTL-biotin-conjugated (1:300, B-1325, Vector Laboratories) and rabbit anti-Jagged1 (1:300, ab7771, Abcam). Confocal imaging was performed using a Zeiss LSM780 scanning confocal, with a Zeiss Plan-Apochromat 25x, 0.8NA multi-immersion objective. Confocal stacks were taken at 1.5 micron Z spacing and exported to Imaris (Biplane) for 3D reconstruction and surface rendering. All other image processing was performed in Fiji (Schindelin et al, Nature Methods, PMID: 22743772). All immunofluorescence analyses were successfully repeated more than three times and representative images are shown.

### QPCR of organoids

Total RNA was extracted from cells using Purelink RNA mini kit (Life Technologies) and cDNA was synthesized from 100 ng total RNA using GoScript™ reverse transcriptase (Promega Australia). qRT–PCR analyses were performed with GoTaq qPCR Master Mix (Promega) by an ABI PRISM 7500 96 real-time PCR machine. All absolute data were first normalized to GAPDH and then normalized to control samples (DDCt method). The sequences of primers used for qRT–PCR are as listed in Supplementary Table 9.

### Isolation of EpCAM-enriched cellular fractions

Kidney organoids were transferred to a drop of trypsin-EDTA, minced with a surgical blade, transferred to a 15 ml tube with 3 ml trypsin-EDTA and incubated at 37 °C for 10 min while gently pipetting the disintegrating organoid pieces every 3 minutes. Next 8 ml of DMEM + 10%FBS was added and cells pelleted by centrifugation (250g 5 min). The pellet was next resuspended in 2 ml of epithelial cell culture medium (Renal Epithelial cell growth medium, cat#C-26030, Banksia Scientific). Remaining cell clumps were removed with a 40 μM cell strainer (BD biosciences) and cells were counted. Cells (10^7^) were repelleted by centrifugation (250g 5 min), resuspended in 200 μl of MACS buffer (MACS buffer: 46.7ml PBS+; 3.3ml of 7.5% BSA and 200 μl of 0.5M EDTA) mixed with 20 μl of CD326 microbeads (Miltenyi Biotec, cat#130-061-101) and refrigerated to 4 °C. After 30 min 5 ml of MACS buffer was added, and cells were pelleted by centrifugation, resuspended in 500 μl MACS buffer and applied to a pre-rinsed MS column in the MACS separator. After 3 washings with MACS buffer cells were collected with 1 ml of MACS buffer after removing the column from the MACS separator. This routinely yielded approximately 3 x 10^5^ EpCAM+ cells/organoid and live cell rates of >75%.

### Isolation of LTL-enriched cellular fractions

Pooled iPSC-derived kidney organoids were dissociated by incubation with TrypLE select enzyme (Thermo Fisher) for 12 minutes at 37°C, with gentle pipetting every 2 minutes to aid dissociation. The cell solution was passed through a series of cell strainers with sequentially smaller mesh sizes, ranging from 100μM to 40μM (Corning) with extensive washing using sterile chilled MACS buffer (D-PBS, 2nM EDTA, 0.5% bovine serum albumin) to yield a single cell suspension. Cell were counted, centrifuged at 300g for 5 minutes and resuspended in 100μl of MACS buffer with 1μl of biotin-conjugated LTL primary antibody (Vector Labs), and incubated on ice for 30mins. Cells were rinsed with MACS buffer and centrifuged twice before resuspending in 300μl of MACS buffer plus 100μl Streptavidin microbeads (Miltenyi Biotech) for 30 mins on ice. Cells were rinsed with MACS buffer and centrifuged before resuspending in 500μL of MACS buffer and passed through an MS MACS column according to the manufacturer’s protocol (Miltenyi Biotec). The LTL+ fraction was eluted from the column, the sorted cell population counted and stored at -80°C until RNA extraction was performed.

## List of abbreviations

iPSC: induced pluripotent stem cells
RNAseq: RNA sequencing

## Declarations

### Ethics approval and consent to participate

Ethics approval for the derivation of human induced pluripotent stem cells line RG_0019.0149.C6 was approved as a subproposal of HREC_14_QRBW_34 by The Human Research Ethics Committee of the Royal Womens and Childrens Hospital.

### Consent for publication

Not applicable

### Availability of data and material

The datasets generated and analysed during the current study are available in the NCBI Gene Expression Omnibus (Edgar et al, 2002), under accession numbers GSE99942, GSE70101, GSE89044, GSE99468, GSE99469, GSE99582 and GSE107305.

### Authors’ contributions

BP advised on experimental design, performed all statistical analysis and wrote the manuscript. PXE, MT, LH, JS and DY performed differentiation experiments. PXE prepared RNA and analysed QPCR data. DY and KL collected and presented morphological IF data. JS and LH performed EPCAM+ and LTL+ MACS sorting respectively. EW and JS generated iPSC cell lines. AO advised on experimental design and oversaw the statistical analysis. ML devised the study, deigned and interpreted all experimental data and wrote the manuscript. All authors read and approved the final manuscript.

## Acknowledgements

We thank Angelika Christ and Greg Baillie at the Institute for Molecular Bioscience, The University of Queensland, for sequencing services. We acknowledge Dr. Andrew Mallett and Dr. Stephen Alexander for assistance in ethics applications and the patient recruitment.

## Supplementary Figures

**Supplementary Figure 1:**
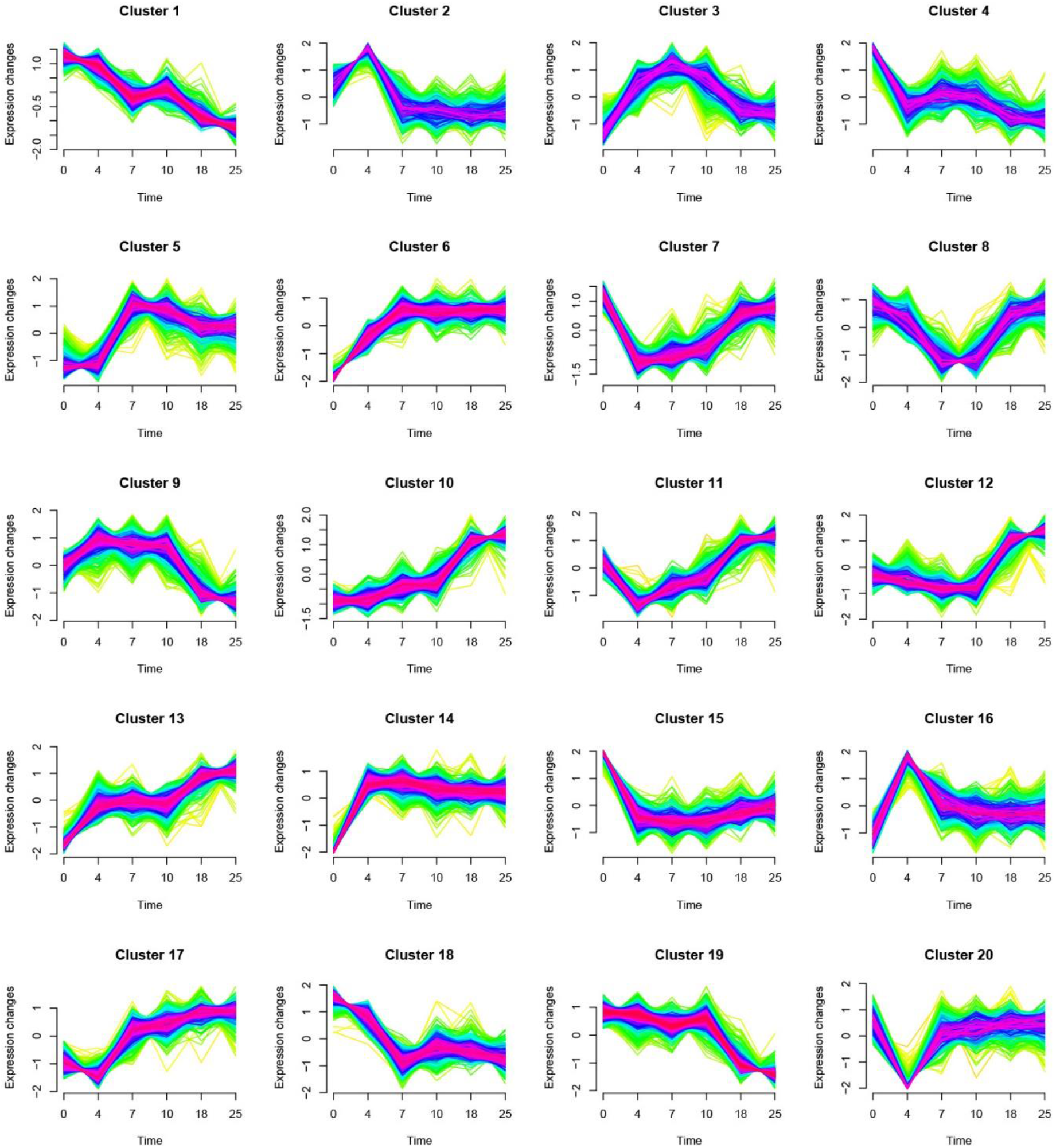
Expression patterns for the 20 fuzzy clusters identified across the time points of the organoid differentiation protocol.

**Supplementary Figure 2:**
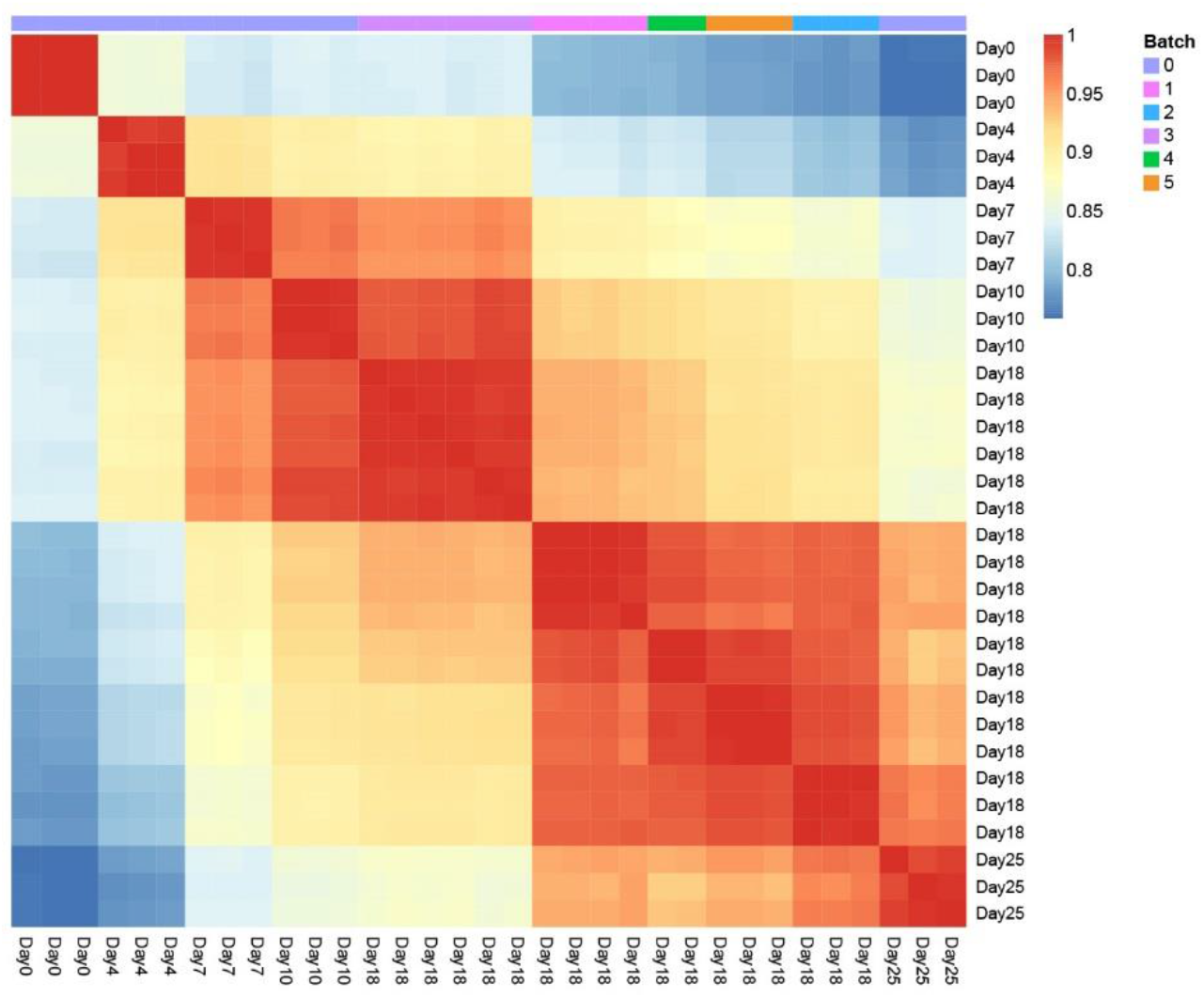
Pairwise Spearman’s correlation between all CRL1502-C32 organoids.

**Supplementary Figure 3:**
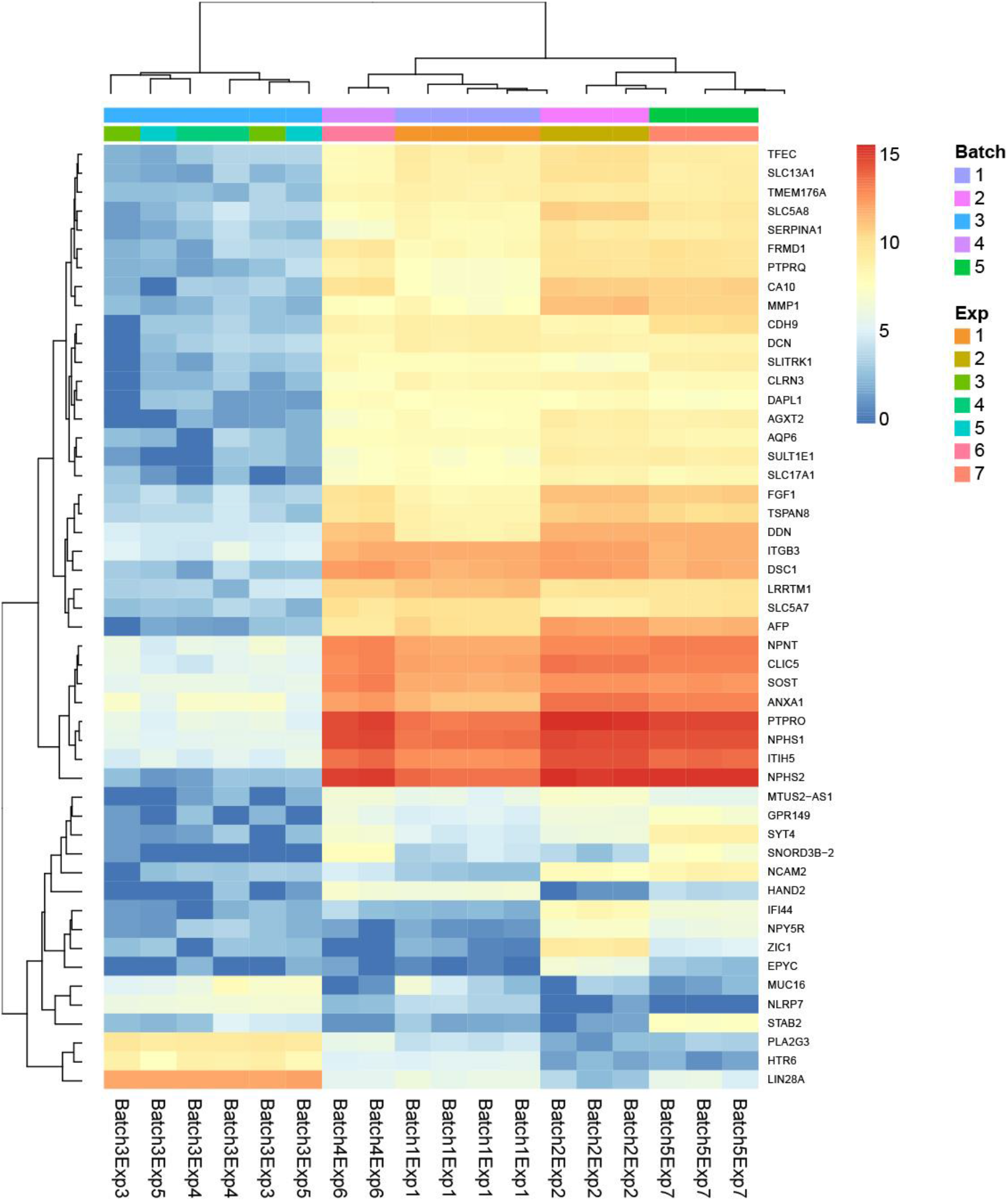
Heatmap for top 50 most variable genes between CRL1502-C32 day 18 organoids. Unsupervised hierarchical clustering shows that organoids within batches cluster together.

**Supplementary Figure 4:**
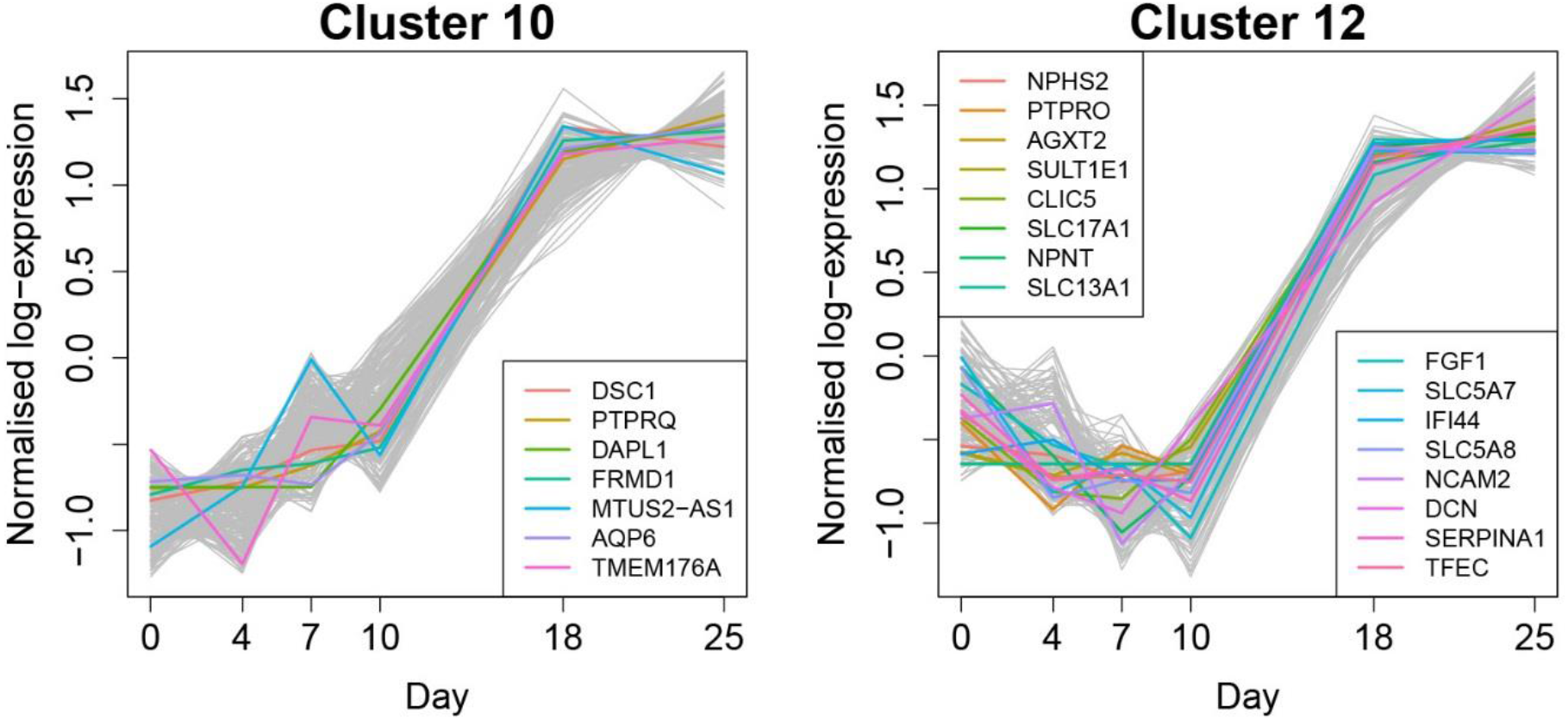
Highly variable genes present in clusters 10 and 12. The overlap between the top 50 most highly variable genes and core genes in each cluster showed 7 and 16 genes in common in clusters 10 and 12 respectively.

**Supplementary Figure 5:**
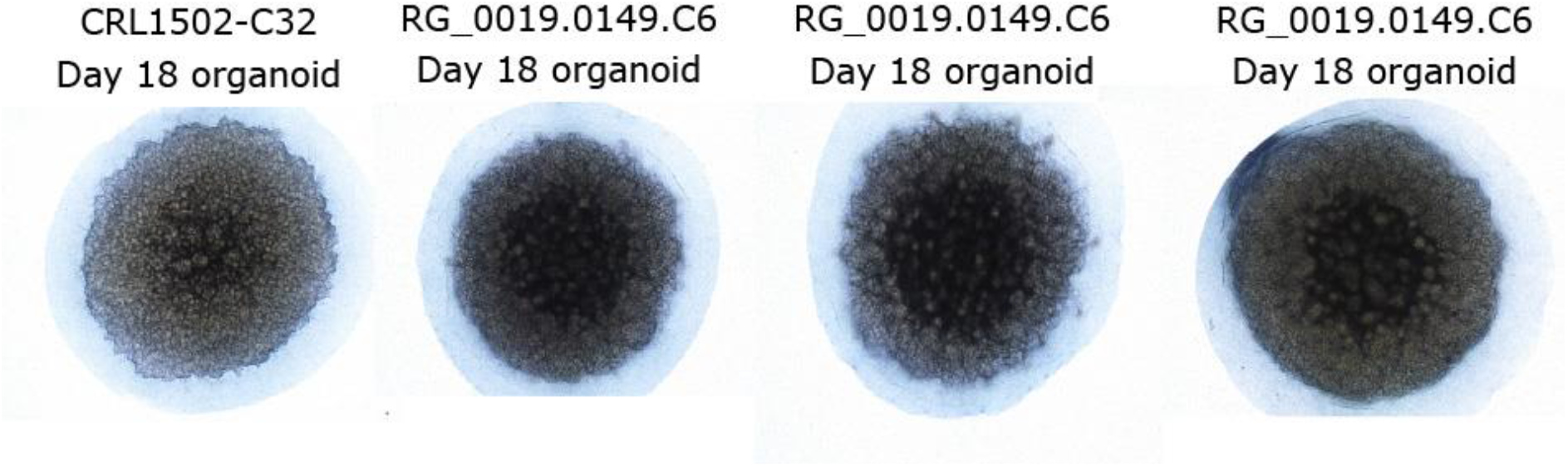
Immunofluorescence of cell line RG_0019.0149.C6 displaying similar nephron formation and segmentation compared to the CRL1502-C32 line.

**Supplementary Figure 6:**
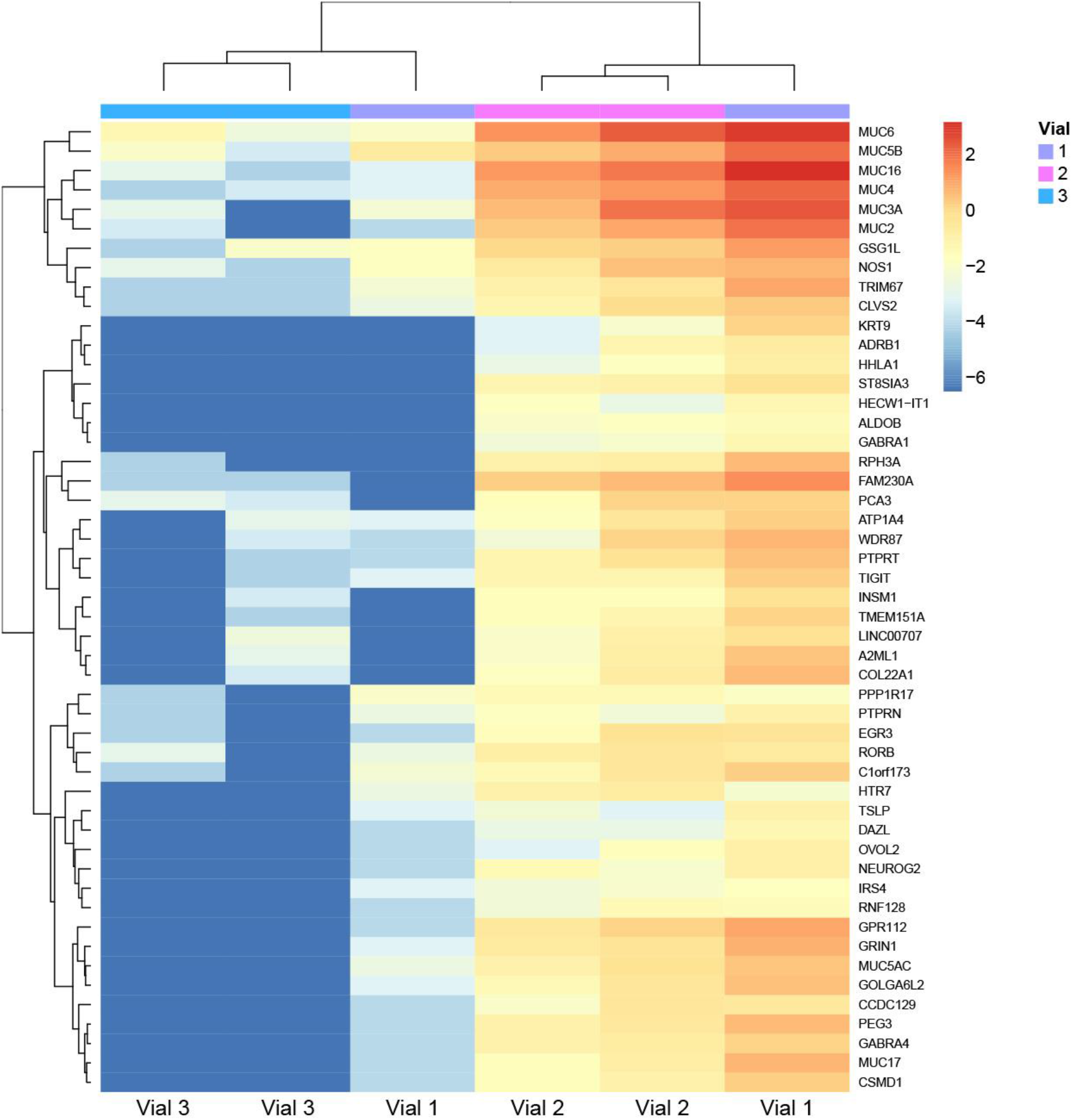
Heatmap for top 50 most highly variable genes between RG_0019.0149.C6 day 18 organoids.

**Supplementary Figure 7:**
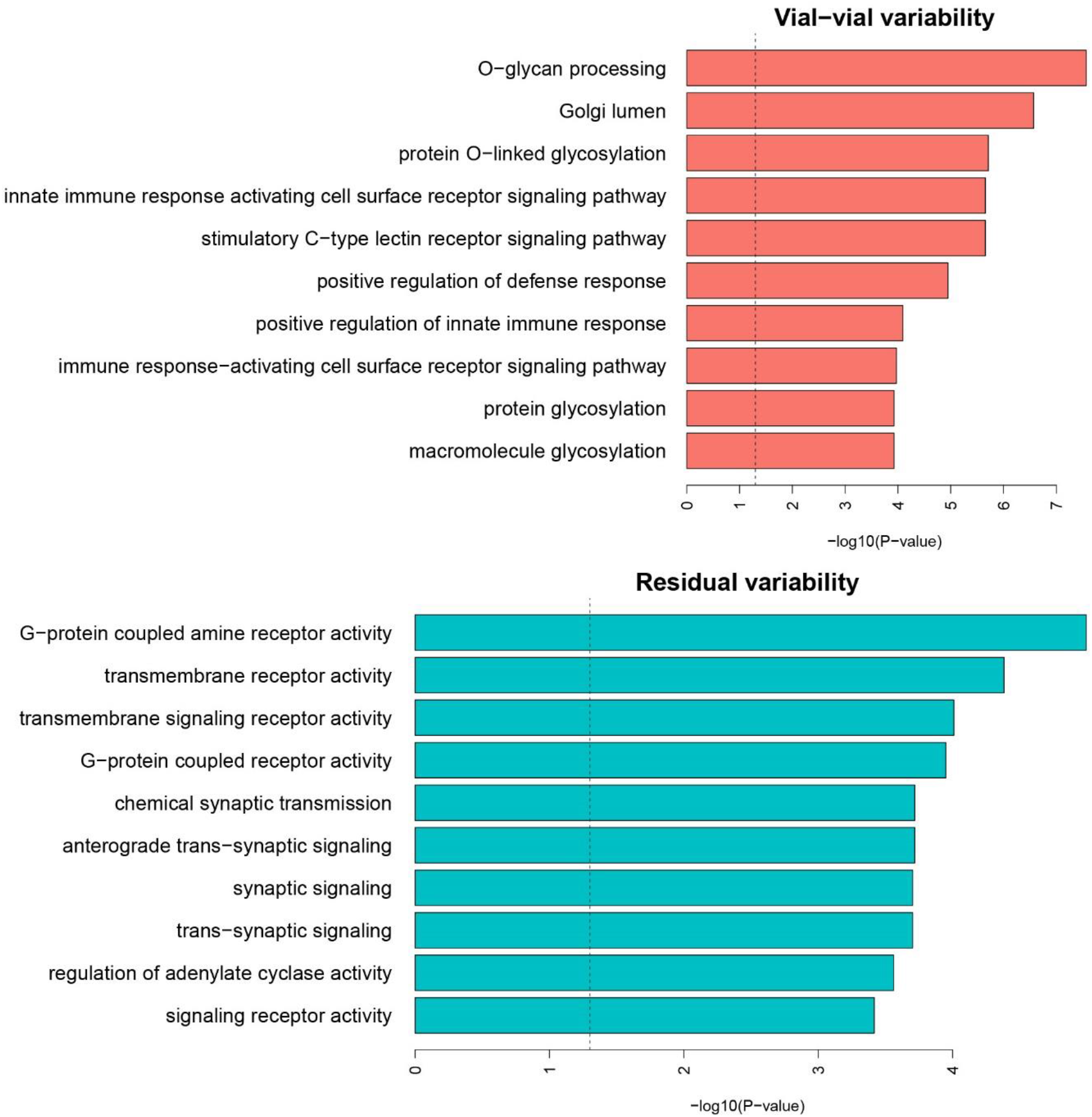
Enriched GO terms for genes contributing to vial-to-vial and residual variability. Top 100 genes contributing to each source of variability were tested.

## Supplementary Tables

**Supplementary Table 1:** Differentially expressed genes between consecutive timepoints for the CRL1502-C32 time series data.

**Supplementary Table 2:** Top 20 enriched GO and KEGG terms for top 100 up and down-regulated genes between consecutive timepoints

**Supplementary Table 3:** Core genes in 20 fuzzy clusters

**Supplementary Table 4:** Top 20 enriched GO and KEGG for core genes in 20 fuzzy clusters

**Supplementary Table 5:** FANTOM5 Transcription Factors (hg19) present in each cluster

**Supplementary Table 6:** Random effects analysis identifying genes that contribute major sources of variation between CRL1502-C32 organoids

**Supplementary Table 7:** Random effects analysis for cell line RG_0019.0149.C6 investigating contributions to vial-to-vial and residual variability

**Supplementary Table 8:** Top 20 enriched GO terms for top 100 genes contributing to vial-to-vial and residual variability for RG_0019.0149.C6

**Supplementary Table 9:** Primers for qRT-PCR

## References

Ader, M. & Tanaka, E.M., 2014. Modeling human development in 3D culture. Current Opinion in Cell Biology, 31, pp.23–28.

Aksoy, I. et al., 2017. Personalized genome sequencing coupled with iPSC technology identifies GTDC1 as a gene involved in neurodevelopmental disorders. Human Molecular Genetics, 26(2), pp.367–382.

Ardhanareeswaran, K. et al., 2017. Human induced pluripotent stem cells for modelling neurodevelopmental disorders. Nature Reviews Neurology, 13(5), pp.265–278.

Bates, D. et al., 2015. Fitting Linear Mixed-Effects Models Using lme4. Journal of Statistical Software, 67(1), pp.1–48.

Bellin, M. et al., 2013. Isogenic human pluripotent stem cell pairs reveal the role of a KCNH2 mutation in long-QT syndrome. The EMBO journal, 32(24), pp.3161–75.

Bouchard, M. et al., 2002. Nephric lineage specification by Pax2 and Pax8. Genes & development, 16(22), pp.2958–70.

Briggs, J.A. et al., 2013. Integration-Free Induced Pluripotent Stem Cells Model Genetic and Neural Developmental Features of Down Syndrome Etiology. STEM CELLS, 31(3), pp.467–478.

Brunskill, E.W. et al., 2008. Atlas of Gene Expression in the Developing Kidney at Microanatomic Resolution. Developmental Cell, 15(5), pp.781–791.

Brunskill, E.W. et al., 2011. Defining the Molecular Character of the Developing and Adult Kidney Podocyte K. Stadler, ed. PLoS ONE, 6(9), p.e24640.

Chen, J. et al., 2009. ToppGene Suite for gene list enrichment analysis and candidate gene prioritization. Nucleic acids research, 37(Web Server issue), pp.W305–11.

Dobin, A. et al., 2013. STAR: ultrafast universal RNA-seq aligner. Bioinformatics (Oxford, England), 29(1), pp.15–21.

Edgar, R., Domrachev, M., Lash, A.E. (2002) Gene Expression Omnibus: NCBI gene expression and hybridization array data repository. Nucleic Acids Res. 30(1),207–10.

Eiraku, M. et al., 2011. Self-organizing optic-cup morphogenesis in three-dimensional culture. Nature, 472(7341), pp.51–56.

Fujimura, N. et al., 2007. Wnt-mediated Down-regulation of Sp1 Target Genes by a Transcriptional Repressor Sp5. Journal of Biological Chemistry, 282(2), pp.1225–1237.

Futschik, M.E. & Carlisle, B., 2005. Noise-robust soft clustering of gene expression time-course data. Journal of bioinformatics and computational biology, 3(4), pp.965–88.

Gentleman, R.C. et al., 2004. Bioconductor: open software development for computational biology and bioinformatics. Genome biology, 5(10), p.R80.

Georgas, K. et al., 2009. Analysis of early nephron patterning reveals a role for distal RV proliferation in fusion to the ureteric tip via a cap mesenchyme-derived connecting segment. Developmental Biology, 332(2), pp.273–286.

Georgas, K. et al., 2008. Use of dual section mRNA in situ hybridisation/immunohistochemistry to clarify gene expression patterns during the early stages of nephron development in the embryo and in the mature nephron of the adult mouse kidney. Histochemistry and Cell Biology, 130(5), pp.927–942.

Harding, S.D. et al., 2011. The GUDMAP database--an online resource for genitourinary research. Development (Cambridge, England), 138(13), pp.2845–53.

Howden, S.E., Thomson, J.A. & Little, M.H., 2017. Simultaneous Reprogramming and Gene Editing of Human Fibroblasts. In Press, Nature Protocols.

Huch, M. & Koo, B.-K., 2015. Modeling mouse and human development using organoid cultures. Development (Cambridge, England), 142(18), pp.3113–25.

Jang, Y.-Y. & Ye, Z., 2016. Gene correction in patient-specific iPSCs for therapy development and disease modeling. Human Genetics, 135(9), pp.1041–1058.

Kadoshima, T. et al., 2013. Self-organization of axial polarity, inside-out layer pattern, and species-specific progenitor dynamics in human ES cell-derived neocortex. Proceedings of the National Academy of Sciences of the United States of America, 110(50), pp.20284–9.

Kim, C. et al., 2013. Studying arrhythmogenic right ventricular dysplasia with patient-specific iPSCs. Nature, 494(7435), pp.105–110.

Lancaster, M.A. et al., 2013. Cerebral organoids model human brain development and microcephaly. Nature, 501(7467), pp.373–379.

Liao, Y., Smyth, G.K. & Shi, W., 2014. featureCounts: an efficient general purpose program for assigning sequence reads to genomic features. Bioinformatics (Oxford, England), 30(7), pp.923–30.

Lindström, N.O. et al., 2015. Integrated β-catenin, BMP, PTEN, and Notch signalling patterns the nephron. eLife, 3, p.e04000.

Little, M.H. et al., 2007. A high-resolution anatomical ontology of the developing murine genitourinary tract. Gene Expression Patterns, 7(6), pp.680–699.

McCarthy, D.J. & Smyth, G.K., 2009. Testing significance relative to a fold-change threshold is a TREAT. Bioinformatics, 25(6), pp.765–771.

McCracken, K.W. et al., 2014. Modelling human development and disease in pluripotent stem-cell-derived gastric organoids. Nature, 516(7531), pp.400–404.

Nakano, T. et al., 2012. Self-Formation of Optic Cups and Storable Stratified Neural Retina from Human ESCs. Cell Stem Cell, 10(6), pp.771–785.

O’Brien, L.L. et al., 2016. Differential regulation of mouse and human nephron progenitors by the Six family of transcriptional regulators. Development (Cambridge, England), 143(4), pp.595–608.

Paquet, D. et al., 2016. Efficient introduction of specific homozygous and heterozygous mutations using CRISPR/Cas9. Nature, 533(7601), pp.125–129.

Phelan, D.G. et al., 2016. ALPK3-deficient cardiomyocytes generated from patient-derived induced pluripotent stem cells and mutant human embryonic stem cells display abnormal calcium handling and establish that ALPK3 deficiency underlies familial cardiomyopathy. European Heart Journal, 37(33), pp.2586–2590.

Risso, D. et al., 2014. Normalization of RNA-seq data using factor analysis of control genes or samples. Nature Biotechnology, 32(9), pp.896–902.

Ritchie, M.E. et al., 2015. limma powers differential expression analyses for RNA-sequencing and microarray studies. Nucleic Acids Research, p.gkv007-.

Robinson, M.D., McCarthy, D.J. & Smyth, G.K., 2010. edgeR: a Bioconductor package for differential expression analysis of digital gene expression data. Bioinformatics, 26(1), pp.139–140.

Robinson, M.D. & Oshlack, A., 2010. A scaling normalization method for differential expression analysis of RNA-seq data. Genome Biology, 11(3), p.R25.

Spence, J.R. et al., 2011. Directed differentiation of human pluripotent stem cells into intestinal tissue in vitro. Nature, 470(7332), pp.105–109.

Suga, H. et al., 2011. Self-formation of functional adenohypophysis in three-dimensional culture. Nature, 480(7375), pp.57–62.

Takahashi, K. et al., 2007. Induction of Pluripotent Stem Cells from Adult Human Fibroblasts by Defined Factors. Cell, 131(5), pp.861–872.

Takasato, M. et al., 2016. Generation of kidney organoids from human pluripotent stem cells. Nature Protocols, 11(9), pp.1681–1692.

Takasato, M. et al., 2015. Kidney organoids from human iPS cells contain multiple lineages and model human nephrogenesis. Nature, 526(7574), pp.564–568.

Tan, G. & Lenhard, B., 2016. TFBSTools: an R/bioconductor package for transcription factor binding site analysis. Bioinformatics, 32(10), pp.1555–1556.

Tsang, T.E. et al., 2000. Lim1 Activity Is Required for Intermediate Mesoderm Differentiation in the Mouse Embryo. Developmental Biology, 223(1), pp.77–90.

Wang, Q. et al., 2005. Odd-skipped related 1 (Odd1) is an essential regulator of heart and urogenital development. Developmental Biology, 288(2), pp.582–594.

Young, M.D. et al., 2010. Gene ontology analysis for RNA-seq: accounting for selection bias. Genome biology, 11(2), p.R14.

Yu J, Hu, K, Smuga-Otto, K et al. Human induced pluripotent stem cells free of vector and transgene sequences. Science 2009; 324: 797–801.

